# Pan-Cancer PDOs Preserve Tumor Heterogeneity and Uncover Therapeutic Vulnerabilities

**DOI:** 10.1101/2025.04.10.647635

**Authors:** Hui-Hsuan Kuo, Bhavneet Bhinder, Hamza Numan Gokozan, Kathryn Gorski, Pooja Chandra, Jyothi Manohar, Daniela Guevara, John Otilano, Jenna Moyer, Marvel Tranquille, Sarah Ackermann, Jared Capuano, Cynthia Cheung, Thomas Anthony Caiazza, Phoebe L. Reuben, Anastasia Murray Tsomides, Adriana Irizarry, Michael Sigouros, David Wilkes, Abigail King, Troy Kane, Majd Al Assaad, Wael Al Zoughbi, Kentaro Ohara, Joonghoon Auh, Peter Waltman, Florencia P. Madorsky Rowdo, Enrique Podaza, Valerie Gallegos, John Nguyen, Raehash Shah, Manish Shah, Allyson Ocean, Douglas Scherr, Nasser Altorki, Melissa Frey, Ana M. Molina, Lisa Newman, Vivan Bea, Eloise Chapman-Davis, Marcus D. Goncalves, Ashish Saxena, Parul J. Shukla, Kevin Holcomb, Rachel Simmons, Scott Tagawa, Jonathan H. Zippin, Evelyn Cantillo, Rohit Chandwani, Melissa Davis, Kelly Garrett, Pashtoon Murtaza Kasi, Jennifer Marti, David Nanus, Jones Trevor Nauseef, Elizabeth Popa, Joseph Ruggiero, Momin T. Siddiqui, Alicia Alonso, Cora N. Sternberg, Bishoy M. Faltas, Olivier Elemento, Juan Miguel Mosquera, Andrea Sboner, M. Laura Martin

**Author notes:** Co-first authors (Hui-Hsuan Kuo, Bhavneet Bhinder). Co-senior authors (Juan Miguel Mosquera, Andrea Sboner, M. Laura Martin). Corresponding authors: M. Laura Martin.

## Abstract

We developed a tumor-matched, pan-cancer patient-derived organoid (PDO) platform comprising 220 PDOs from 190 patients across 15 cancer types to advance functional precision oncology. Our comprehensively characterized PDOs showed 93% histopathology concordance, 80% median genomic concordance for driver mutations, and a 0.85 median gene expression correlation with parent tumors. Gene expression in PDOs remained stable across ≥ 10 passages, supporting reproducibility for long-term drug screening. Even PDOs with low genomic concordance retained oncogenic drivers, supporting their use as disease models. Clonality analysis revealed that 85% of PDOs preserved dominant tumor clones. Higher genomic concordance was associated with greater clonal similarity, while lower genomic concordance was associated with clonal divergence. Functional assays showed that 58% of PDOs from a subset of patients ineligible for FDA-approved PARP inhibitors responded to Talazoparib, with sensitivity linked to alterations in DNA damage repair. Combination screens revealed drugs that effectively overcame resistance, especially in TP53-mutant PDOs. In summary, our platform supports investigation of targeted therapies, identification of molecular features linked to drug sensitivity, and translational discovery, offering insights into personalized cancer treatment beyond current biomarker guidelines.

## INTRODUCTION

Functional precision oncology platforms enable direct testing of cancer therapies in patient-derived tumor tissues, identifying vulnerabilities that may not be detectable through molecular profiling alone^1,2^. This approach, crucial for advancing precision medicine, addresses the complexity and heterogeneity of tumors, offering treatment options to patients without actionable biomarkers or with acquired drug resistance. Within functional precision oncology, patient-derived organoids (PDOs) have emerged as promising pre-clinical models. PDOs are increasingly integrated into functional precision oncology, alongside advanced technologies like next-generation sequencing and clinical data, to uncover patient-specific molecular features and guide the development of more effective, targeted therapies.

PDOs have been generated from a wide variety of cancer types and pan-cancer PDO cohorts have been reported^3,4^. These renewable resources are invaluable to study disease progression, test drug efficacy, conduct toxicity studies, and may even serve as patient avatars for treatment and co-clinical trials^5–7^. To ensure reliable results from these models, it is important to standardize and validate the PDO platforms used for such applications. However, inconsistencies among existing PDO pipelines, particularly in model characterization and validation — meaning the extent to which PDOs recapitulate key molecular, histological, and functional features of the parent tumor — remain a significant challenge. For example, criteria to determine successful generation of PDOs are not universally defined, strategies to evaluate their fidelity to parent tumors at molecular level are not routinely applied, and biomarker assessment for cancer subtype classification are not consistently performed. Additionally, concerted efforts to collect clinical data for screened PDOs are lacking, even though such information is crucial to correlate drug responses with patient characteristics. As PDO pipelines continue to evolve and integrate into functional precision oncology, robust standardization efforts are necessary to ensure their effectiveness for personalized drug testing and reliability as models to predict patient responses to therapies.

To support this goal, we present a comprehensively characterized pan-cancer PDO platform comprising 220 PDOs from 190 patients, validated against parent tumors for histopathology, genomic features, and expression profiles. This work advances our PDO platform introduced in 2017, broadening its characterization and application^3^. Here, we report high rates of concordance between PDOs and parent tumors in tissue and cytomorphology^8,9^, somatic mutations, somatic copy number alterations (SCNAs), and expression profiles, confirming the platform’s reliability to generate pre-clinical models that faithfully recapitulate their matching tumor biology. As a proof of concept for its application in functional precision oncology, we tested the sensitivity of a subset of PDOs to Talazoparib, a poly (ADP-ribose) polymerase inhibitor (PARPi). These PDOs were derived from cancer patients who, based on current FDA guidelines, were not eligible for PARPi-based therapies^10–14^. This application highlighted the capability of our platform to investigate targeted therapies and identify genomic and transcriptomic trends associated with drug sensitivity, while proposing potential combination therapy strategies across a wide range of cancers. Our well-characterized PDO platform offers promising pre-clinical models to advance precision oncology research with a goal to improve patient care.

## RESULTS

### Development and Characterization of Successful Patient-Derived Organoid Models

We developed a robust PDO platform to establish, characterize, and leverage PDOs from tissues across 15 different cancer types. Between 2014 and 2022, we received 1,472 tissue samples and initiated cultures for 1,408. We observed successful PDO formation in 81% (n = 1,140/1,408) of samples (Fig. 1A). Cultures that failed to form PDOs at passage 0 (p0) were either contaminated with bacteria, particularly those from research autopsies^15^, or did not have enough viable cells. PDOs that did not grow beyond five passages (p5) or failed to expand sufficiently for biobanking were classified as short-term cultures (53.5%, 610/1,140) (Fig. 1A). We successfully biobanked 128 (11.2%) cultures that showed robust expansion to 1×10^6^ cells for cryopreservation before reaching p5, with an additional 402 cultures (35.3%) continuing to grow and were biobanked after p5. (Fig. 1A). PDOs that expanded consistently to p5 were characterized by their histopathology and molecular features and compared to their parent tissue (Fig. 1A). Those characterized by histopathology review and/or molecular profiles were classified as established PDOs. Based on these criteria at the time of data freeze in mid-2023, we present here a pan-cancer cohort of 220 established PDOs from 191 patients (Fig. 1A-B, Table 1).

**Figure 1.**
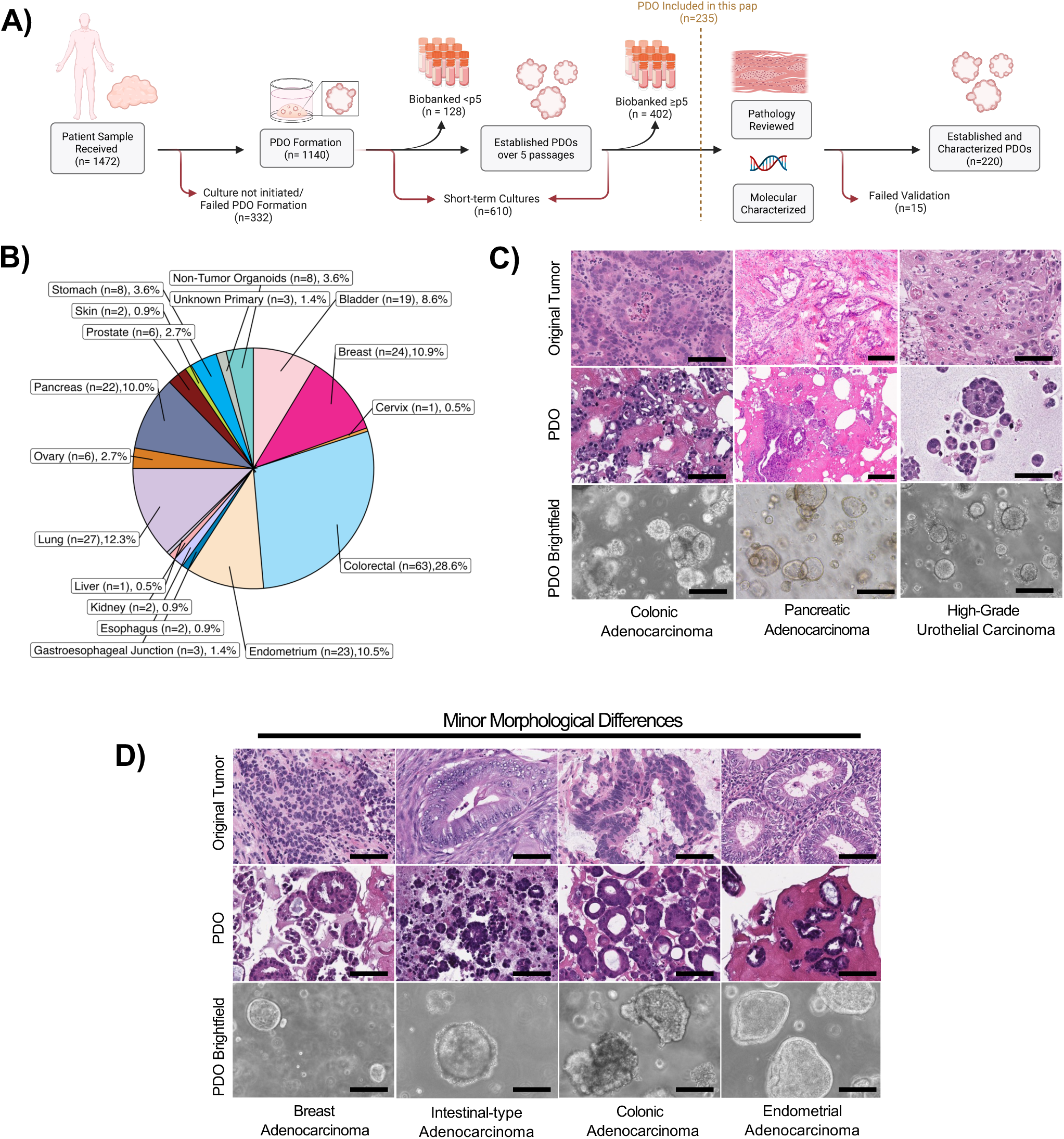
Overview of the PDOs pan cancer cohort. **A)** PDO development and characterization workflow. **B)** Pie chart showing the number and fraction of established PDOs by primary tumor type. **C)** H&E-stained slides of three representative pairs of PDOs and parent tumors which have highly concordant tissue morphologies. **D)** H&E-stained slides of four representative PDO and parent tumor pairs, illustrating minor morphological differences. Left: A breast adenocarcinoma PDO exhibiting tubular and cribriform structures, in contrast to the solid growth pattern of the parent tumor. Middle Left: A gastrointestinal adenocarcinoma PDO with papillary/micropapillary architecture and a high nuclear-to-cytoplasmic ratio, compared to the rounded tubular configuration with columnar nuclei in the parent tumor. Middle Right: A colon adenocarcinoma PDO with rounder nuclei, contrasting with the distinctly columnar nuclei of the parent tumor. Right: An endometrial adenocarcinoma PDO showing a relatively higher nuclear-to-cytoplasmic ratio than the parent tumor. Scale bar: 150 μm.

**Table 1.** Patient and sample characteristics for established PDOs.

| Characteristics | PDOs, Total N (%) |
| --- | --- |
| **Cohort** |  |
| Patients | 191 |
| Samples | 220 |
| **Age** |  |
| Median (range) years | 63 (23-94) |
| **Sex Assigned at Birth** |  |
| Female | 80 (42%) |
| Male | 74 (39%) |
| Not Available | 37 (19%) |
| **Self-reported Race** |  |
| Asian or Pacific Islander | 9 (5%) |
| Black | 23 (12%) |
| White | 98 (51%) |
| Not Listed | 10 (5%) |
| Declined | 12 (6%) |
| Not Available | 39 (21%) |
| **Self-reported Ethnicity** |  |
| Hispanic or Latino | 8 (4%) |
| Non-Hispanic | 113 (59%) |
| Declined | 20 (11%) |
| Not Available | 50 (26%) |
| **Prior Treatment** |  |
| No pretreatment | 18 (9%) |
| 1-3 pretreatment | 79 (42%) |
| 3+ pretreatment | 43 (23%) |
| No information | 51 (27%) |

The success rate of PDO formation with our workflow improved significantly from 57.1% to 81.8% over time and varied by sample source, with higher formation fluid samples (92.5%) compared to autopsy samples (57.6%), as well as cancer type, with the highest success in endometrial cancers (94.2%) and the lowest rates in prostate cancer (69.4%) (Supplementary Fig. 1A-C). PDOs for some tumor types required longer time to successfully establish than others, probably due to suboptimal growth conditions or inherently slower tumor proliferation. For example, the median time to establish breast PDOs was 86.5 days, compared to 36 days for ovarian tumors (Supplementary Table 1).

In our cohort, bladder, breast, endometrium, and lung cancer PDOs were mostly derived from primary lesions (over 80%), while colorectal, pancreas, skin, and prostate cancer PDOs were predominantly metastatic (over 65%) (Supplementary Table 1). Colorectal (28.6%), lung (12.3%), endometrium (10.5%), breast (10.9%), and pancreatic (10%) cancers represented the most common tumor types in the established cohort (Fig. 1B). Our PDO collection also included a small subset of melanomas (0.9%), cancers of unknown primary (1.4%) and normal organoids, *i.e*., PDOs from benign tissues (3.6%) (Fig. 1B).

In summary, our cohort of 220 PDOs, established from 191 patients from multiple race groups across 15 different cancer types and benign tissues with well-characterized molecular features, offers a diverse and robust set of models for functional precision oncology research. Table 1 summarizes the cohort demographics, including biological sex, self-reported race and ethnicity, as well as total number of treatments received prior to tissue collections.

### Patient-Derived Organoids are Morphologically Similar to Parent Tumors

We performed a thorough histopathology review of hematoxylin and eosin (H&E) stained sections of 198 PDOs and their parental tissues to determine if tumor cytomorphology and cellular characteristics were maintained in the organoids^8,9^. A select subset of immunohistochemistry (IHC) markers were used to confirm biomarker expression.

A high concordance was observed in 93% (185/198) of PDOs closely matching their parent tissues. Most of them (78%; 154/198) mirrored the cellular architectures of their parental tumors and were consistent with the histopathology of the given tumor type. For example, PDOs derived from colon and pancreatic adenocarcinomas preserved their characteristic tubular (glandular) structures. Similarly, PDOs from high-grade urothelial carcinomas retained features of the parent tumors, including evidence of divergent differentiation, such as squamous morphology (Fig. 1C). The remaining subset of PDOs, 31 out of 198 (15.7%), showed minor discrepancies where the PDO was morphologically slightly different than the parental tumors with either different cytomorphology (21/31), histology grade (7/31) or biomarker expression (3/31) (Fig. 1D, Supplementary Table 2). Only 7% (13/198) of the samples failed pathological review potentially due to contaminations of the tissue samples with neighboring cells, resulting in normal cell overrepresentation from cancerous tissue (8/13), benign-appearing tissues that yielded tumor cell growth (4/13), or a potential mix up of two PDO samples (1/13) (Supplementary Fig. 1D, Supplementary Table 2). In some cases, small clones — either normal or cancerous — expanded to become the dominant population in organoid culture, further emphasizing the need for rigorous pathological review. To address these issues two histopathologists independently assessed the neighboring sections to determine the degree of contamination in our samples and, if needed, recharacterized the PDOs as either cancerous or normal. These measures to refine the classification of our PDOs ensured their reliability and accuracy as disease models for further research.

### Genomic Features Are Conserved between PDOs and Parent Tumors

The next step in the pipeline was to assess how closely PDOs represented their parent tumors by comparing their genomic features. We analyzed mutational profiles of 147 unique pairs of PDOs and parent tumors using Whole Exome Sequencing (WES: n = 114 pairs) and targeted gene panels (n = 33 pairs) (Supplementary Table 3). We identified a strong positive correlation among tumor-PDO pairs for tumor mutation burden (TMB) (r = 0.79, p < 2.2e-16) and microsatellite instability (MSI) scores (r = 0.94, p < 2.2e-16) (Fig. 2A). The ploidy correlation was modest (r = 0.34, p < 0.00025); however, after excluding 21 outlier pairs (18% of the dataset), the correlation significantly improved to 0.81 (Supplementary Fig. 2A-B).

**Figure 2.**
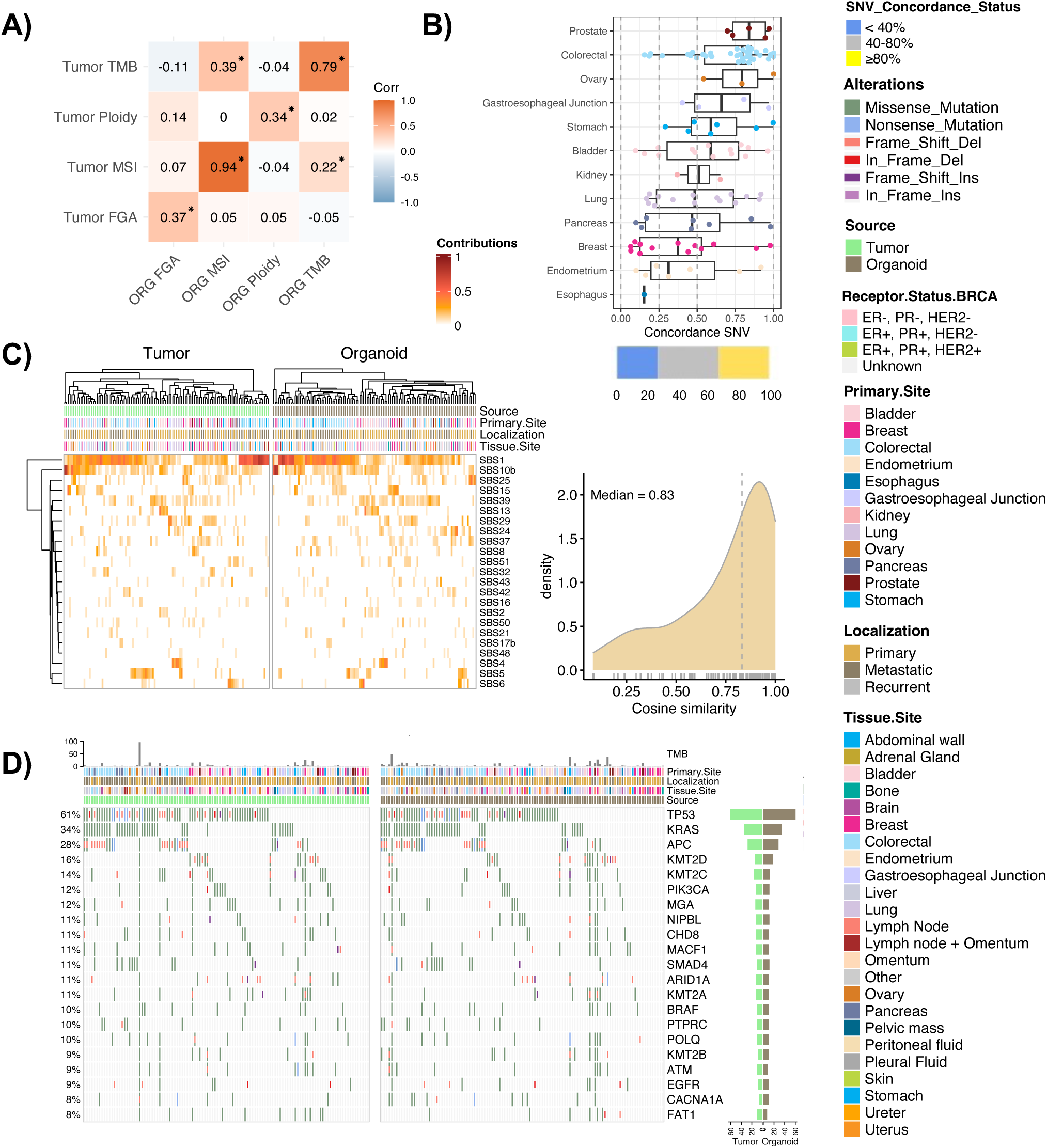
Comparative analysis of genomic characteristics and mutation profiles between PDOs and parent tumors. **A)** Pearson correlation among PDO and parent tumors for aggregate measures of genomic characteristics. Asterisk (*) denotes a significant correlation (p-value < 0.05). FGA, fraction genome altered; MSI, microsatellite instability; TMB, tumor mutation burden. **B)** Boxplots showing the distribution of genomic concordance between PDOs and parent tumors across 12 cancer types, calculated based on the proportion of shared non-silent mutations. Bar plot at the bottom shows the percentage of samples that belong to the low (< 40%), mid (40-80%) and high (≥ 80%) concordance groups. **C)** Heatmap showing the most frequent COSMIC SBS mutational signatures, selected based on contributions > 0 in a minimum number of samples in the cohort. The density distribution plot on the right shows the distribution of the cosine similarity scores of COSMIC SBS signature profiles between paired PDOs and parent tumors. **D)** Oncoprint displaying the top 21 most frequent cancer driver genes in the cohort, selected based on a mutation frequency of 9% or higher in either the tumor or the PDO sample groups. Bar plots on the right indicate the frequency of each mutated gene within the tumor and PDO sample groups.

To assess genomic fidelity at the mutation level between tumors and their matched PDOs, we calculated a metric we termed somatic nucleotide variant (SNV) concordance, defined here to include single nucleotide variants and small insertions/deletions. This metric represented the proportion of tumor mutations also detected in the matched PDOs. Across the cohort, the median SNV concordance was 60%, with approximately one-third of PDO-tumor pairs showing high SNV concordance (≥ 80%) (Fig. 2B). SNV concordance varied by cancer type, being highest for prostate tumors (84%, n = 5) and lowest for the only esophagus tumor in the cohort (15%, n = 1) (Fig. 2B)^16,17^. The low concordance in this single esophageal pair could be attributed to the low inferred cancer cell purity of 27% for the PDO. But, unexpectedly, in general the SNV concordance was independent of tumor purity in our cohort (Pearson r = 0.13, p-value = 0.18) as for the pairs with SNV concordance of ≤ 40%, the average purity was 92.5% in PDOs and 66% in tumors (Supplementary Table 3, Supplementary Fig. 2C).

When we restricted our WES SNV concordance analysis to 200 pan-cancer driver genes^18^, our SNV concordance increased to 80% (Supplementary Fig. 2D, Supplemental Table 3). These results are in alignment with the previously reported somatic concordance of 77.6% for a pan-cancer cohort^4^. For targeted panels, our SNV concordance (median = 88%) was even more strongly conserved, with 15 out of 33 tumor-PDO pairs showing a 100% concordance (Supplementary Fig. 2E-F, Supplemental Table 3). Consistent with previous reports, these findings confirm a robust SNV concordance between PDOs and parent tumors in our cohort, especially for known cancer driver genes.

In addition to SNV concordance, we tested if the characteristic patterns of somatic mutations were conserved between PDOs and parent tumors. We quantified the contributions of 65 COSMIC mutation signatures for each sample and identified strongly conserved mutational signature profiles between the PDO and parent tumor pairs (cosine similarity = 0.83)^19,20^ (Fig. 2C). Single base substitution 1 (SBS1), the clock-like aging signature, was the most common signature among the samples in our cohort, consistent with reports of SBS1 being the most common COSMIC signature found across many cancer types^21^. We also identified conserved signatures that have known prognostic associations with survival in certain cancer types. For example, the APOBEC3-induced mutational signatures SBS2 and SBS13 were highly conserved between the bladder cancer PDO-tumor pairs (cosine similarity = 0.97)^22^ suggesting that PDOs faithfully recapitulate this important mutational process.

### Patient-Derived Organoids Recapitulate Driver Mutations of Parent Tumors

Cancer driver mutations are critical for selecting models in drug screening, especially for targeted therapies and precision medicine. To evaluate our models, we assessed whether PDOs exhibited any bias toward specific driver mutations. Our analysis revealed no significant differences in driver mutation rates between PDOs and their corresponding parent tumors (Fig. 2D, Supplementary Table 4). This finding remained consistent when the analysis was extended to include all mutated genes across the cohort (FDR = 1) (Supplementary Table 5).

To further examine how well PDOs recapitulated individual driver mutations, we compared driver mutation frequencies in PDOs to those observed in parent tumors, and to expected frequencies in corresponding cancer types from The Cancer Genome Atlas (TCGA) (Fig. 2D, Supplementary Fig. 2F). We identified *TP53* as the most frequently mutated gene in our cohort (tumor = 61%; PDO = 60%; n = 114), consistent with its status as the most frequently mutated gene in TCGA pan-cancer cohorts (44.6%, n = 6,408). Besides *TP5*3, *KRAS* (tumor = 34%; PDO = 34%), and *APC* (tumor = 28%; PDO = 28%) were the top mutated genes, a pattern representative of the higher proportion of colorectal tumors in our cohort (∼31%) (Fig. 2D, Supplementary Table 4).

To test if PDOs served as good disease-specific models, we analyzed the frequency of cancer-specific mutations within each corresponding cancer type (Supplementary Fig. 3A-H, Supplementary Table 6). For most PDOs, driver mutations overlapped well with their corresponding parent tumors and occurred at expected frequencies. For example, colorectal tumors (n = 34) were enriched in *TP53* (tumor = 74.3%; PDO = 73.5%), *KRAS* (tumor = 65.7%; PDO = 67.6%), and *APC* (tumor = 57.1%; PDO = 55.9%) mutations (Supplementary Fig. 3A). These genes were also the top three most frequently mutated drivers in the TCGA-colon adenocarcinoma (COAD) and TCGA-rectum adenocarcinoma (READ) cohorts (*TP53* = 59.5%, *KRAS* = 42.2%, and *APC =* 74.2%)^23,24^. Similarly, lung tumors (n = 15) were enriched in *KRAS* (Tumor = 46.7%, PDO = 46.7%, TCGA-lung adenocarcinoma (LUAD) = 46.1%), *TP53* (Tumor = 40%, PDO = 46.7%, TCGA-LUAD = 50.8%), *EGFR* (Tumor = 20%, PDO = 26.7%, TCGA-LUAD = 14.3%), and *KEAP1* (Tumor = 20%, PDO = 20%, TCGA-LUAD = 17.4%) mutations (Supplementary Fig. 3B)^23,24^. Bladder tumors (n = 14) were enriched in mutations for *TP53* (Tumor = 64.3%, PDO = 64.3%, TCGA-bladder urothelial carcinoma (BLCA) = 49.2%), *KMT2D* (Tumor = 50%, PDO = 50.0%, TCGA-BLCA = 27.7%), *PIK3CA* (Tumor = 35.7%, PDO = 28.6%, TCGA-BLCA = 20%), and *FGFR3* (Tumor = 28.6%, PDO = 21.4%, TCGA-BLCA = 12.3%) (Supplementary Fig. 3C). These findings show that PDOs faithfully recapitulate the driver mutation frequencies of their parent tumors and closely mirror TCGA cohort data, supporting their utility as robust models for disease-specific drug screening and precision medicine.

### Clonal Architecture Partly Explains Differences in Genomic Concordance

As noted above, most PDOs retained cancer specific driver mutations, including those with low SNV concordance (< 40%), supporting their relevance as good experimental models (Supplementary Fig. 3A-H, Supplementary Table 7). In breast cancer, however, low-concordance pairs (n = 13) lacked key driver mutations (Fig. 3A, Supplementary Table 7), raising the possibility that perhaps clonal selection in PDO cultures may have contributed to these discrepancies. To investigate this, we inferred the clonal composition of all breast cancer samples in a paired tumor-organoid (TO) analysis (Fig. 3B, Supplementary Fig. 4A). In a representative low-concordance pair (WCM2108, Concordance = 6%), the tumor and PDO showed distinct subclonal populations, whereas in a high-concordance pair (WCM2137, Concordance = 98%) tumor and PDO showed substantial overlap in shared clones (Fig. 3B). However, this pattern was not observed uniformly across all cases (Supplementary Fig. 4A), suggesting that while clonal selection may contribute to discordant mutation profiles, it did not fully explain them.

**Figure 3.**
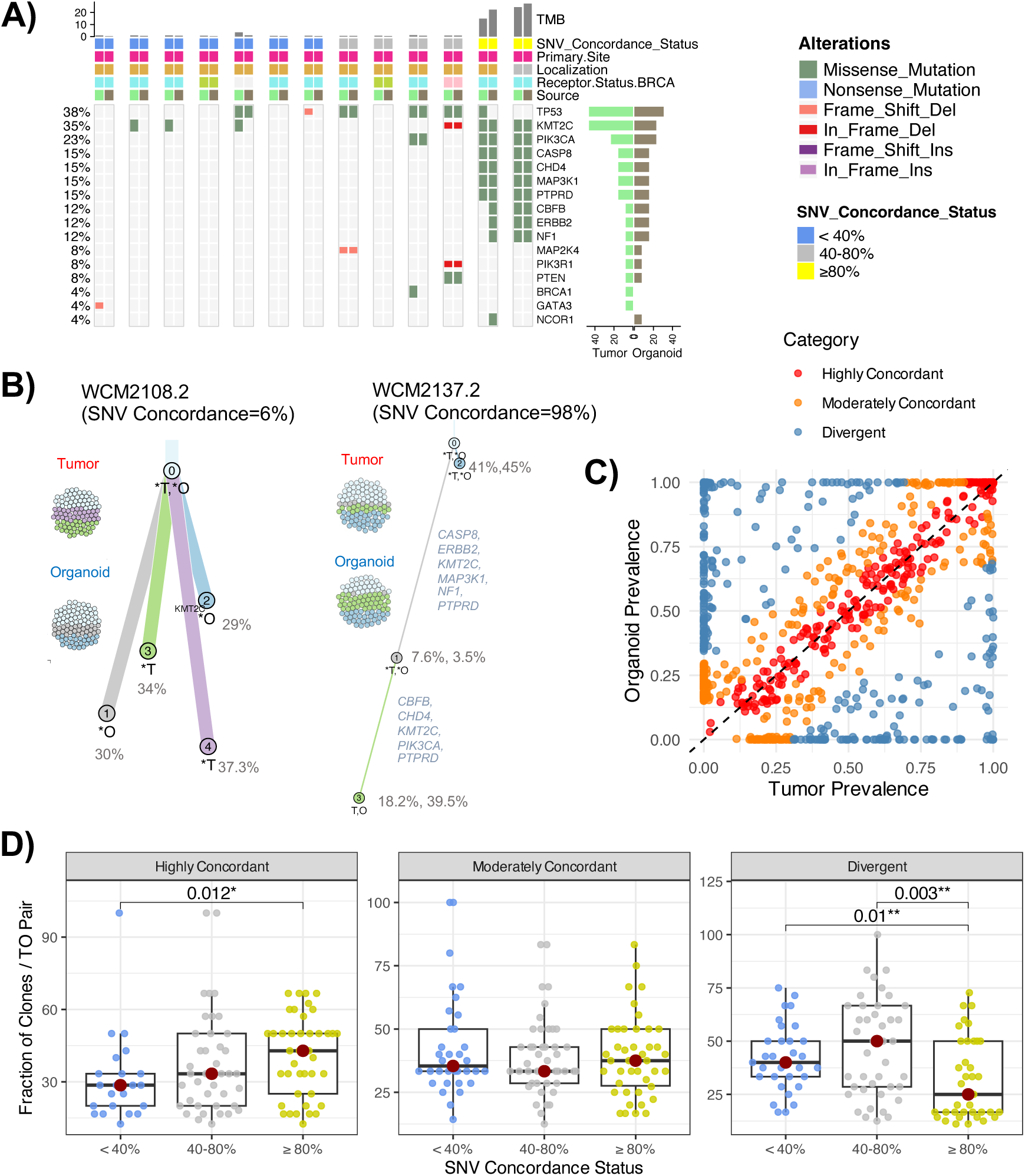
Cellular prevalence of mutations in PDOs compared to matched tumors. **A)** Oncoprint showing the frequency of breast cancer-specific driver genes^18^ in a paired analysis of breast tumors and PDOs, ordered by genomic concordance levels: < 40%, 40-80%, and ≥ 80%. **B)** Graphical representation of shared clonal architecture between pairs of PDOs and parent tumors, featuring one with the lowest concordance (left) and another with the highest concordance (right) from the breast cancer sub-cohort. Clonal trees are also annotated with known driver mutations in the corresponding samples. Sphere of cells to the left of each tree represent proportion of clonal subpopulations in a sample out of a 100%. **C)** Scatter plot comparing cellular prevalence of mutation clusters inferred using PyClone-VI. Each point represents a distinct cluster, with its estimated prevalence in the tumor (x-axis) and the corresponding PDO (y-axis). Points are colored by degree of concordance. The diagonal line represents equal prevalence in both PDO and matched tumor**. D)** Boxplots displaying the distribution of the fraction of clones per TO pair within each clonal divergence category (Highly Concordant, Moderately Concordant, Divergent), stratified by SNV concordance status (< 40%, 40–80%, ≥ 80%). Each dot represents one TO pair in that divergence category. Median values of each boxplot are marked by dark red dot. P values are from pairwise Wilcoxon tests and only significant p values < 0.05 are shown. Clone classification was based on absolute differences in cellular prevalence inferred using PyClone-VI.

To better understand global patterns of clonal preservation and divergence across tumor types, we extended our analysis to the full cohort by applying PyClone-VI to all available TO pairs (n = 132)^25^. Since PyClone-VI clusters mutations with similar cellular prevalence under the assumption that they arise from the same cellular lineage, we interpreted each mutation cluster as representing a distinct clonal population. We then assessed clonal divergence by comparing the cellular prevalence of each cluster between tumor and PDO. Clusters were categorized as Highly Concordant (<10% absolute difference), Moderately Concordant (10 to 29%), or Divergent (≥ 30%) based on their absolute difference in cellular prevalence. Remarkably, based on the absolute difference in cellular prevalence between tumor and PDO, we found that the majority of clones were either highly concordant or moderately concordant (62.2%), indicating substantial preservation of clonal structure (Figure 3C). When overlaid with SNV concordance levels, TO pairs with high SNV concordance (≥ 80%) showed significantly higher fraction of highly concordant clones (p = 0.01) and a lower fraction of highly divergent clones (p = 0.01) compared to those with low SNV concordance (< 40%) (Figure 3D). These findings suggested that genomic similarity is associated with clonal stability during organoid derivation.

The patterns observed also pointed to possible expansion, contraction, or loss of clones during PDO establishment and growth. To quantify these dynamics, we calculated the fold change in cellular prevalence between tumor and PDO for dominant tumor clones (defined as those with > 10% prevalence in the tumor). At the cohort level, 85% of dominant tumor clones were retained in the matched PDO, while 15% (n = 96) were lost. Among retained clones, 24% (n = 153) showed contraction (fold change < 0.75) and 18% (n = 116) showed expansion (fold change > 1.25). We further stratified these patterns by tumor type and observed that, across most cancer types, a substantial fraction of dominant clones remained stable (Unchanged) or increased in prevalence (Expanded) in PDOs, supporting the overall preservation of key clonal populations (Supplementary Fig. 4B). Notably, colorectal and bladder tumors displayed the highest median proportion of Unchanged clones (approx. 67%), whereas stomach tumors showed broader variability in clonal outcomes, including elevated rates of clonal loss and expansion. These findings point toward tissue-specific variability in how clonal populations are reshaped in PDO cultures.

### Divergence in CNA Profiles between PDOs and Parent Tumors

To determine the genomic fidelity of established PDOs for CNAs, we performed a comparative analysis similar to the one conducted for mutations. We calculated CNA concordance based on the number of CNA events shared between PDOs and parent tumors relative to the total number of events in the parent tumor (Fig. 4A). Our pan-cancer cohort had a median CNA concordance rate of 74% and nearly half of these pairs had a high concordance of ≥ 80% (Fig. 4A). These concordance rates accurately captured the shared patterns of genomic CNAs between the PDO-tumor pairs (Fig. 4B). CNA concordance was marginally correlated with SNV concordance per PDO-tumor pair (r = 0.26, p-value = 0.0076, n = 105) (Fig. 4C). Similar to mutations, CNA concordance did not correlate with tumor purity, ploidy, or the fraction of the genome altered. (Fig. 4D, Supplementary Fig. 5A-B).

**Figure 4.**
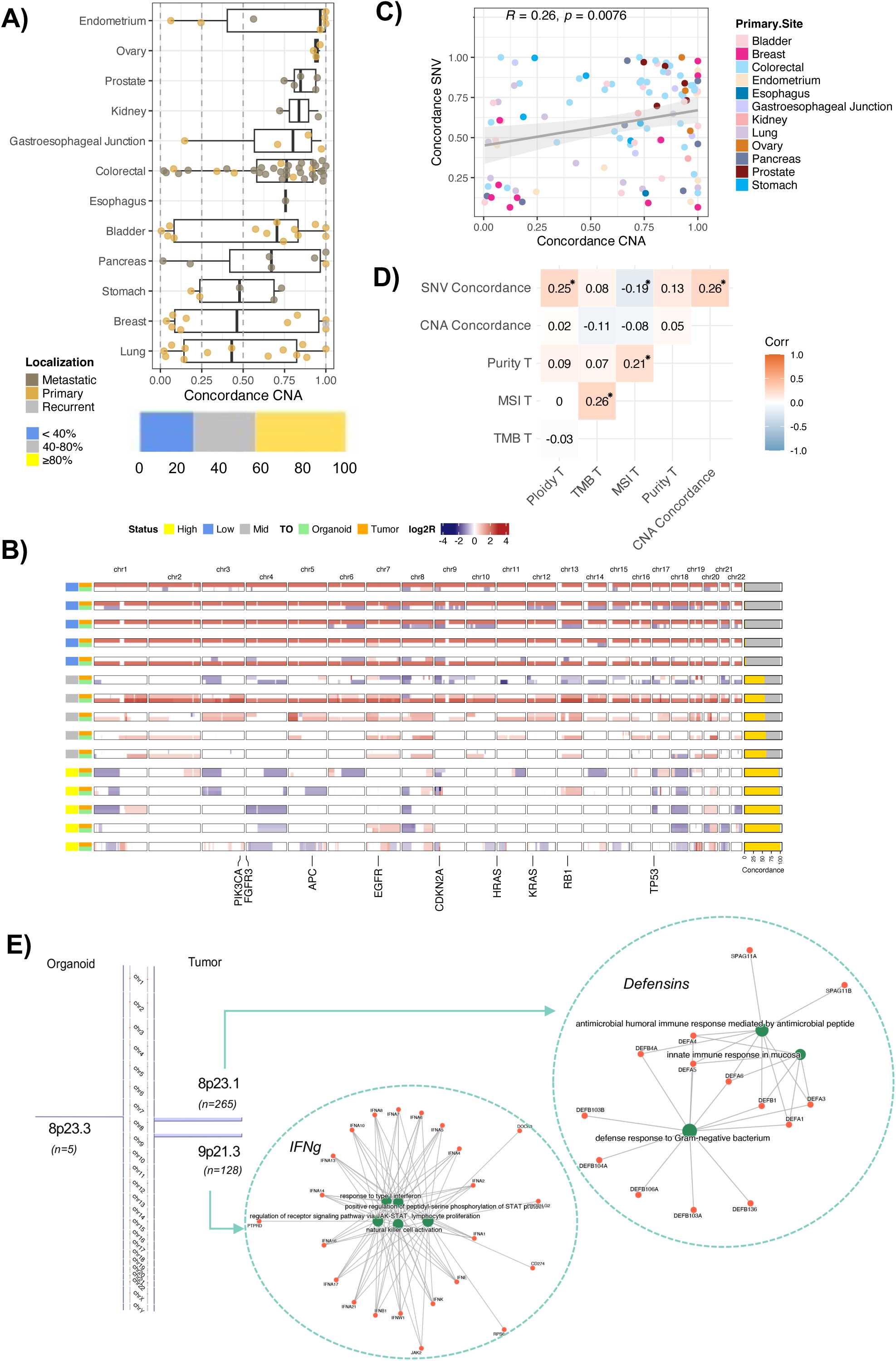
Comparative analysis of CNA between PDOs and parent tumors. **A)** Boxplots showing the distribution of CNA concordance between PDOs and parent tumors across 12 cancer types, calculated based on the proportion of shared amplified, deleted, or neutral regions in the genome. Bar plot at the bottom shows the percentage of samples which belonged to the low (< 40%), mid (40-80%) and high (≥ 80%) CNA concordance groups. **B)** Heatmap showing the log2 ratios for the copy number segments across the human exome (excluding chromosomes X and Y) in a paired PDO-tumor analysis. The heatmap shows five representative samples each from the low (< 40%), mid (40-80%) and high (≥ 80%) CNA concordance groups. **C)** Scatter plot with a linear regression line (and a gray region for 95% confidence intervals) between the CNA (x-axis) and mutation/SNV concordance (y-axis) for the cohort. The points are colored by cancer type. **D)** Pearson correlation among the two types of concordance quantified and the purity, MSI and TMB of the parent. Asterisk (*) denotes a significant correlation (p-value <0.05). **E)** The significantly deleted copy number peaks (blue) identified in the GISTIC analysis of the tumor samples (right) or in the PDOs (left). The number of genes in the peak are indicated as n in the brackets under the cytoband descriptor. The networks denote the enriched biological processes associated with the genes in the cytobands (as marked by the blue arrows and circles).

In our CNA concordance analysis, we identified 28 PDO-tumor pairs with a low concordance rate of < 40%. To better understand this low concordance, we further investigated these pairs for known driver CNAs, similar to our approach with mutations. We observed that PDOs with < 40% concordance had poor overlap in driver CNAs with their parent tumors. However, driver CNAs were still present in the PDOs, even though they did not exactly match those in the parent tumors (Supplementary Fig. 5C). This finding motivated us to collect PDOs with low concordance (< 40% with parent tumors) for both CNA and SNV (n = 12) and characterize them for driver events independent of their parent tumors. In this analysis, we identified key cancer drivers (such as *TP53*, *KRAS*) in PDOs (Supplementary Fig. 5D), indicating that despite their low overall concordance with parent tumors, these PDO still harbor essential cancer-driving alterations and could be good standalone cancer models.

We examined whether certain CNA events were enriched in PDOs relative to tumors in general. We analyzed CNA profiles using Genomic Identification of Significant Targets in Cancer (GISTIC) to identify significant CNAs separately for PDOs and tumors (Supplementary Fig. 6A-B, Supplementary Table 8). Among the large cluster of genes, we identified loci *8p23.1* and *9p21.3*, which were significantly deleted in tumors but not in PDOs (Fig. 4E). The deleted regions of loci *8p23.1* enriched in defensins genes and *9p21.3* enriched in interferon gamma (IFN-γ) genes. Deletion in loci *9p21.3* has previously been shown to be associated with an immune cold tumor microenvironment (TME) and a poor response to immune checkpoint immunotherapy^26^. Tumors experience heightened selection pressure in the TME from immune surveillance, therefore loss of IFN-γ loci might be one of the mechanisms of immune escape for the tumor, whereas PDOs are not subject to the same selective pressure *in vitro*. In summary, our analysis suggests that while PDOs generally maintain a high degree of CNA concordance with their parent tumors, significant divergences in specific CNA events, particularly those associated with immune selection, may occur.

### Transcriptomic Drift and Differences among Biological Processes and Pathways

To quantify the conservation of key biological processes and pathways between the tumors and the early (passage number ≤ 7) and late passage PDOs, we compared the expression profiles collected for these samples. Overall, an unsupervised clustering of the early passage PDO expression profiles showed a cancer specific clustering of the samples, except for ovarian tumors where the ovarian primary (n = 3) and metastatic (n = 1) samples did not cluster together (Fig. 5A). The expression profiles for early PDOs and matched tumor had a good overall concordance (median r = 0.85 [0.72-0.95], n = 72 pairs) for the top 5,000 most variable genes. This concordance was significantly higher than that observed for randomly paired tumor-PDO backgrounds (p-value < 0.001) (Fig. 5B). Similarly, expression profiles for matched early and late passage PDOs had significantly higher concordance (median r = 0.92) compared to randomly paired PDOs (median r = 0.44), indicating preserved gene expression over multiple passages (Fig. 5C).

**Figure 5.**
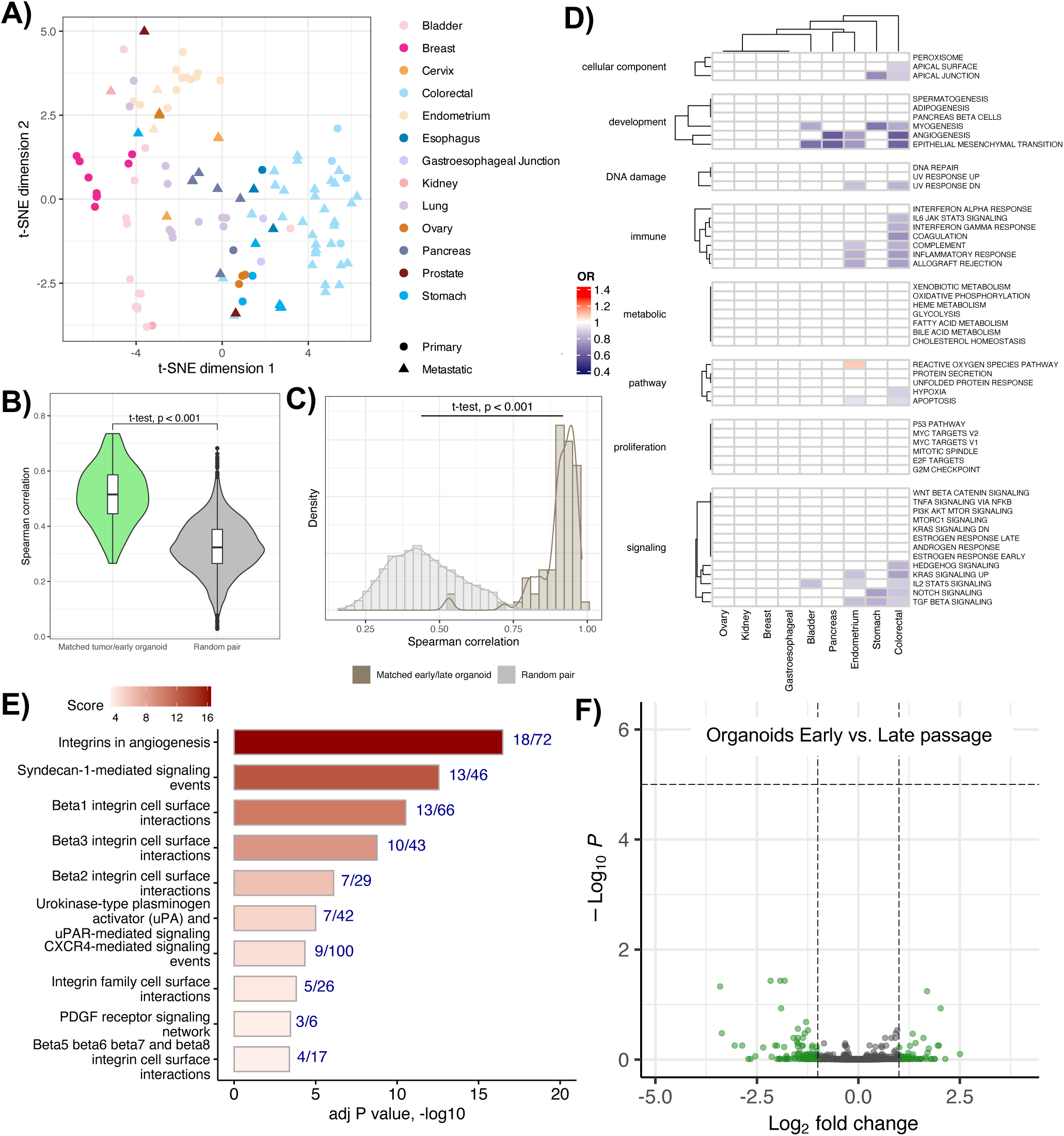
Comparative analysis of gene expression profiles between PDOs and parent tumors. **A)** t-distributed Stochastic Neighbor Embedding (tSNE) plot to show the clustering of PDOs based on their top 5,000 most variable genes. Samples are colored by tumor type and shaped based on their localization. Plot shows cancer types with ≥ 2 PDOs. **B)** Violin plots showing the distribution of Spearman correlation coefficients for top 5000 most variable genes between expression profiles of tumors and PDOs from early passage (passage number ≤ 7) shown in green and between randomly matched pairs shown in gray. **C)** Histograms showing the distribution of correlations between matched early and late passage PDOs against a randomly matched pair background. **D)** Clustered heatmap of differentially regulated 50 cancer hallmark pathways between PDOs and parent tumors. The heatmap shows the odds ratios (OR) of differences in pathway expression levels, where color red denotes higher, and color blue denotes lower pathway expression in PDOs compared to the tumors. Only significant OR (p < 0.01) are shown in the plot. **E)** Barplot showing the enriched pathways in the genes differentially downregulated in at least 5 different cancer types. X-axis denoted the adjusted p-values on a -log10 scale. The barplot is annotated by the number of downregulated genes found in a pathway versus total number of genes in that pathway. **F)** Volcano plot for DGE analysis between early (passage number ≤ 7) and late passage PDOs (passage number > 7) showing no differentially regulated genes between the two groups.

Next, we aimed to understand whether the cancer hallmark pathways differed between the PDOs and parent tumors. We calculated single-sample GSEA (ssGSEA) scores for each pathway in the Cancer Hallmark pathway collection (n = 50) from MSigDB^27^. In cancer-specific models, we applied a linear regression analysis to test the association of parent tumors and early passage PDOs with each Cancer Hallmark pathway ssGSEA scores and reported the exponent of coefficients as odds ratio (OR) (Fig. 5D). We adjusted these models for tumor purity, to account for purity associated difference among the PDOs and tumors. Majority of the hallmark pathways were not differentially enriched in either the tumors or the PDOs. However, we did identify some differences among the colorectal tumors (n = 26 pairs) where the tumors showed enrichment in pathways associated with the TME such as immune responses and angiogenesis, which are not retained in the PDOs (Fig. 5D)^28^. We repeated the same analysis for the early *versus* late passage PDOs and found no significant differences in any of the hallmark pathways in any of the cancer types analyzed, further supporting the stability of the expression profiles.

To quantify the extent of transcriptomic drift between parent tumors and PDOs, we performed differential gene expression analyses between early-passage PDOs and their matched tumors, and between early- and late-passage PDOs, using paired, cancer-specific comparisons. In our analysis of early-stage PDOs versus tumors, we defined pan-cancer differentially expressed genes (DEGs) as those that were consistently upregulated or downregulated in at least 5 out of the 7 cancer types analyzed (Supplementary Fig. 7A-B). With this selection, we identified 241 downregulated and zero upregulated pan-cancer DEGs in early-stage PDOs relative to parent tumors. These downregulated genes showed a significant enrichment in the ß-integrin family of genes that are associated with the composition and organization of the extracellular matrix (Fig. 5E, Supplementary Table 9). The downregulated DEGs included genes involved in extracellular matrix organization, such as collagens (*COL1A1, COL3A1, COL6A3, COL12A1, COL16A1, COL4A1, COL5A1, COL5A2, COL8A1, COL8A2, COL1A2, COL15A1*), proteoglycans, and glycoproteins (*ACAN, BGN, DCN, LUM, SPARC, PRELP, ASPN*). Additionally, the downregulated DEGs had genes related to cell adhesion and migration including cadherins and integrins (*CDH11, CDH5, CDH6, ITGA8, ITGA11, ITGB3, VCAM1*), as well as adhesion/junctional proteins (*JAM3, CLEC14A, PECAM1, CLDN5*). We also found markers associated with angiogenesis and vascular development (*VEGFR1 [FLT1], VEGFR3 [FLT4], PDGFRB, PDGFRA, KDR, ANGPT2, TEK, ENG, TIE1, CD34, CDH5, ESAM, VWF, PECAM1 [CD31], CLDN5, RAMP2*) as well as surface markers for immune cells (*CD2, CD53, CD248, LSP1, ITGAL, ITGAX, ITGB3, PTPRC, MS4A6A, MS4A7*) and cytokine/chemokine signaling (*CSF1R, CSF2RB, FCGR2A, FCGR3A, CXCL12, IFI44L, FCER1G*). Together these findings showed that the early-stage PDOs lack critical features of the TME, resulting in a diminished ability to mimic the complex interactions between the tumor cells and the stromal/immune compartments of the TME^28^.

We also performed a similar DEGs analysis between early passage and late passage PDOs and did not identify any DEGs between the two groups, indicating a conserved expression landscape across multiple passages (Fig. 5F).

### PDO as Functional Precision Medicine Models for PARPi Sensitivity

Having established that our PDO platform reliably recapitulates parent tumor biology, we aimed to demonstrate the application of PDOs as functional precision oncology models using PARPi drug sensitivity screens. We selected PDOs across gastrointestinal (GI), bladder, ovary, and prostate cancers and optimized a 7-day treatment for capturing long-term cellular response and DNA damage effects, as confirmed by imaging and viability assays (Supplementary Fig. 8A). Using a threshold of 75% cell death at 10 μM, PDOs were classified as sensitive (n = 16) or resistance (n = 17) to Talazoparib (Fig. 6A). We evaluated RAD51, pγH2AX, and geminin staining as potential indicators for PARPi sensitivity but found no significant correlation with Talazoparib response (Spearman r = 0.37, p-value = 0.07), primary tumor type, nor prior platinum response (Fisher’s exact test, p-value = 0.26 and 0.24, respectively) (Supplementary Fig. 8B-D)^29,30^.

**Figure 6.**
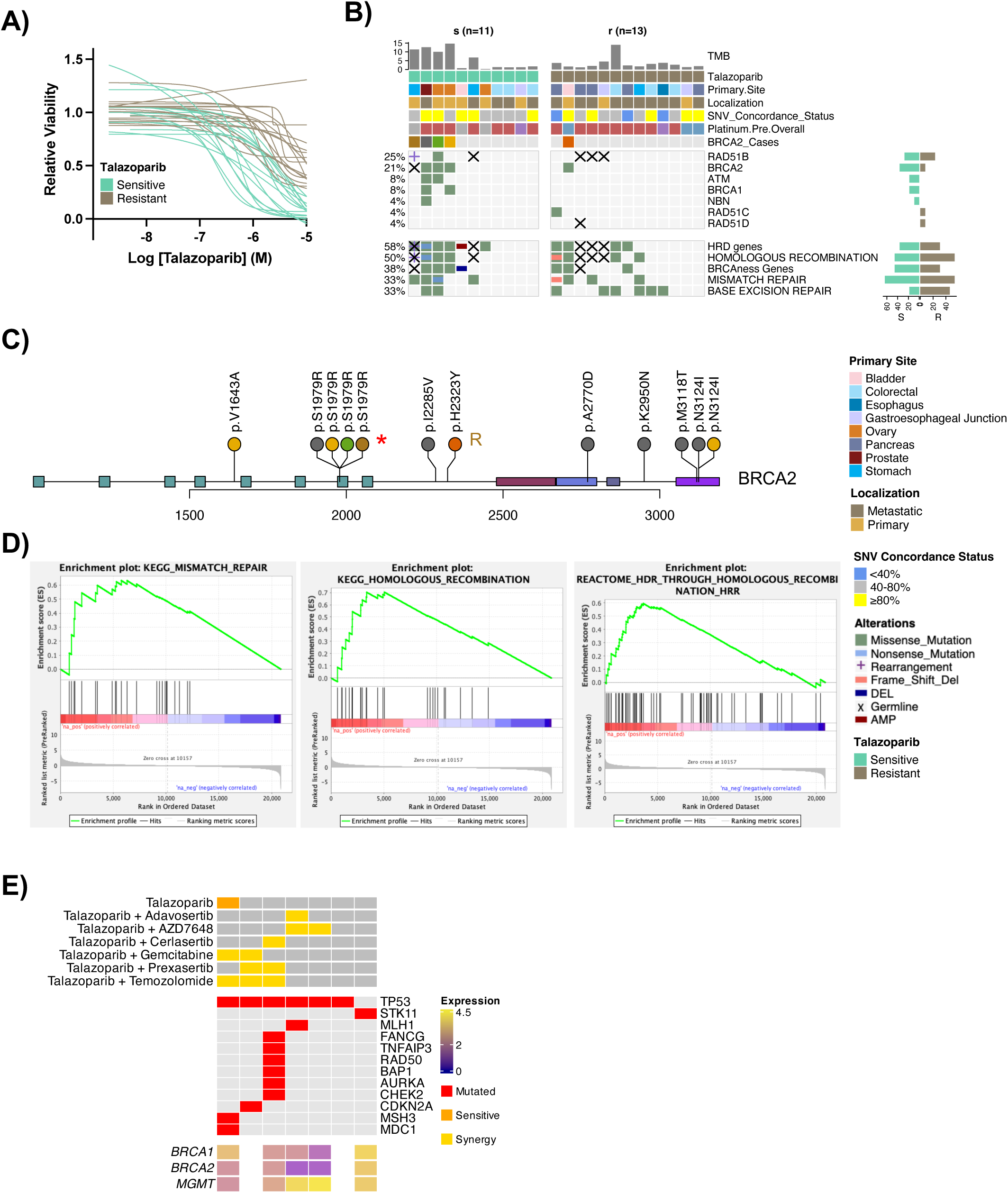
PDOs as a functional precision oncology model to test their sensitivity against Talazoparib. **A)** Dose response curves for Talazoparib sensitivity in PDOs. **B)** Oncoprint of genes relevant to PARPi activity, with PDO samples grouped by sensitivity to Talazoparib. Barplots to the right show gene frequencies in the sensitive (s) versus the resistant (r) PDO groups. HRD stands for Homologous Recombination Deficiency. **C)** Lollipop plot showing variants across PDOs that harbored BRCA2 gene mutations (n = 5). Each color represents a different PDO sample, consistent with the “BRCA2_Cases” annotation bar in the oncoprint shown in panel E. Asterisks denotes germline variant, while an ‘R’ marks the PDO resistant to Talazoparib. **D)** GSEA enrichment plots showing upregulation of mismatch repair and homologous recombination pathways in PDOs sensitive to Talazoparib compared to those resistant to it. Pre-ranked gene list used for this analysis was obtained from the DGE analysis between Talazoparib sensitive versus resistant (reference group) samples post treatment. **E)** Oncoprint showing genetic alterations and expression profiles of selected genes in PDOs treated with Talazoparib, along with their synergy responses to combination treatments with selected inhibitors.

To identify molecular characteristics associated with sensitivity to Talazoparib, we analyzed the mutation profiles of PDOs classified as either sensitive (n = 11) or resistant (n = 13) to the drug, based on sequencing data (Fig. 6B). Among genes relevant to PARPi sensitivity, Talazoparib-sensitive PDOs had a higher frequency of mutations in *BRCA1* (n = 4), *BRCA2* (n = 2), *ATM* (n = 2), and *NBN* (n = 1) genes compared to resistant PDOs. This finding aligns with expectations, as mutations in *BRCA1* and *BRCA2* genes impair DNA repair pathways, rendering cancer cells more sensitive to PARPis. In contrast, mutations in the base excision repair pathway were more common in resistant PDOs (46%) than in sensitive PDOs (18%), which could indicate an adaptive resistance mechanism. Among the *BRCA2*-mutated PDOs, those sensitive to Talazoparib had higher TMB (median TMB: mutated = 12, wild-type = 1.9). Additionally, *BRCA2*-mutated PDOs sensitive to Talazoparib harbored multiple mutations in the *BRCA2* gene, including the pathogenic variant *N3124I*^31,32^. Notably, all Talazoparib-sensitive PDOs carried *BRCA2 S1979R,* a variant of uncertain significance (VUS), in either germline or somatic alterations (Fig. 6C).

Next, we analyzed all nonsynonymous mutations, including those outside DNA repair pathways, to identify those differentially associated with Talazoparib sensitivity. We noted that Talazoparib-sensitive PDOs had mutations in key cellular processes that intersect with DNA repair, including chromatin remodeling (*CREBBP, PHF12*), oxidative (*SLC7A11*) or cellular stress responses (*FMNL1, CRISP3*), DNA repair gene activation (*ALPK2*) and epigenetic regulation (*UHRF1BP1L, ZC3H4*) (Supplementary Fig. 8E).

To investigate differences in expression profiles, we analyzed DEGs between Talazoparib sensitive (n = 14) and resistant (n = 10) PDOs, revealing distinct pathway enrichments at baseline. Talazoparib sensitive PDOs showed upregulation of pathways associated with mismatch repair, homologous recombination deficiency (HRD), extracellular matrix (ECM), and cell adhesion, likely increasing their reliance on PARP-mediated repair (Fig. 6D, Supplementary Fig. 8F). Conversely, downregulation in antigen processing and presentation, PD-1 signaling, and interferon pathways (e.g., interferon gamma) were observed in the Talazoparib sensitive PDOs (Supplementary Fig. 8G). Together with the observed high TMB, these downregulated pathways suggest immune evasion mechanisms possibly at play in the sensitive PDOs.

Finally, we tested combination treatments to sensitize 7 pancreas and GI cancer PDOs with moderate resistance to Talazoparib. We selected 6 drugs targeting key components of the DNA damage repair or previously proposed to have synergistic effect with PARP inhibitors: Adavosertib (Wee1 inhibitor), AZD7648 (DNA-PK inhibitor), Cerlasertib (ATR inhibitor), Gemcitabine (nucleoside analog), Prexasertib (CHK1 inhibitor), and Temozolomide (alkylating agent), with the latter 3 FDA-approved oncology drugs. All 6 drugs enhanced Talazoparib sensitivity to varying degrees (Fig. 6E, Supplementary Fig. 9). The strongest synergy was observed with Temozolomide (synergy scores > 20) in 3 out of 7 PDOs, with 2 additional lines approaching the threshold (8.8 and 9.4). Consistent with our previous findings, all of these lines harbor TP53 alterations^33^. Prexasertib showed synergy in 3 lines, while Gemcitabine, Cerlasertib, and AZD7648 exhibited synergy in 2 lines each. PDOs with moderate baseline sensitivity to Talazoparib responded to more synergistic combinations than resistant PDOs, particularly those with additional DNA repair pathway alterations beyond TP53. Notably, synergy profiles varied across PDOs, highlighting the treatment response heterogeneity and the importance of using PDOs as functional models for personalized combination strategies.

Overall, these findings highlight the complexity of Talazoparib sensitivity, showing that it is not limited to mutations in BRCA genes or prior response to chemotherapy. Instead, multiple factors may influence sensitivity to PARPi. In addition, several DNA damage repair inhibitors exhibited synergy with Talazoparib, though the degree of synergy varied across PDOs based on their genomic composition. These results emphasize the importance of a functional precision oncology approach to identify optimal therapeutic combination for individual patients.

## DISCUSSION

We successfully established and comprehensively characterized a cohort of 220 PDOs from 190 patients and demonstrated their application in functional precision oncology by conducting a PARPi drug screens on a subset. Our cellular prevalence and clonality analysis, spanning multiple tumor types, revealed a level of resolution that provides new insights into the extent and variability of clonal preservation during PDO establishment—an aspect not previously characterized at this scale. This work further validates the robustness of our pan-cancer PDO platform, initially introduced in 2017^3^.

Standardizing PDO generation is key to advancing functional precision oncology. An important consideration in this process is the success rate of PDO cultures, which varies widely depending on tumor type and culture conditions. In this study, the PDO formation rate was approximately 81%, with 46% successfully biobanked, which was an improvement over the 38.6% success rate reported in our initial work^3^. While some studies do not clearly specify their success criteria^34–41^, a few define success based on the ability to expand PDOs^42–44^, whereas others focus on the number of passages or the ability to biobank cultures^3,4,17,41,45–48^. Reported success rates in these cases generally range from 20-70%. Conversely, studies with more stringent criteria that require more than 10 passages, molecular characterization, or long-term cultures exceeding 6 months, tend to report lower success rates of around 10%^17,48^, with an exceptional study achieving a remarkable 70% success rate in establishing neuroendocrine carcinoma organoids^48^. Platforms seeking to establish PDOs as a resource for both functional precision oncology and drug discovery and validation should prioritize models that can expand beyond p5 and can be banked at a minimum of 1×10^6^ cells per vial. Based on these criteria, our platform’s success rates aligned well with other large-scale PDO studies^4^, confirming the robustness of our approach in establishing long-term cultures.

Beyond success rates, validating PDO models against their parent tumors is also essential to ensure accurate representation of the parental tissue. Most studies perform histopathology and genomic profiling to characterize PDOs, but not all report the degree of concordance with the parent tumors^36,47^. Similar to our study, while most studies report good morphological and marker staining concordance, by showing representative images, some have reported contamination with normal cells within their cohorts^34^. Genomic analysis typically involves reporting SNVs and/or CNAs using different sequencing methods, including whole genome sequencing^36,37,42,48^, WES^3,17,35,43,46,47,49,50^, or targeted panels^4,39–41,44,45^. Due to the differences between sequencing methods in depth and coverage, direct comparisons of genomic concordance among different studies are challenging. Targeted panels (100∼500 genes) tend to yield higher concordance^4,40,41,45^ than when broader coverage of ∼20,000 genes, such as WES, is used^3,36,43,47,48^. Studies using autologous pairs of PDOs and tissues generally show good correlations for SNVs and CNAs (60%-90%)^3,4,35–43,47,48^. In cases where parental tissues are unavailable, studies often compare top mutated cancer genes with those in publicly available cohorts^4^. Some studies also investigate genomic stability across passages, typically demonstrating consistent concordance^17,41^. Regarding transcriptomic analysis, most studies examine variations in gene expression under different media conditions^4^, across passages^37,41^, and in relation to IHC or subtype classifications^45–47^. In our cohort, we report a median SNV concordance of 60% (80% when only considering cancer driver genes) and a median CNA concordance of 74% from 114 pan cancer matched pairs using WES, which is comparable to our own previously reported concordance from 14 tumor-PDO pairs^3^. Additionally, our targeted panels demonstrated 88% concordance, which aligns with another pan-cancer platform of 32 tumor-PDO pairs (76.9%)^4^. Overall, our PDOs maintained stable expression profiles between early (p5) and late (p10) passages.

A small subset of our PDOs (∼20%) showed variable degrees of histopathological discrepancies. In cases of minor discrepancies, most PDOs maintained over 50% SNVs concordance, while those with major discordance showed low concordance (0-25%). Despite these discrepancies, PDOs with minor histological discrepancies still showed sufficient genomic concordance to be useful models (Supplementary Table 2). Additionally, 12 PDOs with low concordance (< 40%) for both SNVs and CNAs still retained key cancer driver mutations (Supplementary Table 3). While such deviations from parent tumors might make some PDOs unsuitable for functional precision oncology, their expandability and ability to preserve key cancer-related features highlight their value as cancer disease models to study cancer etiology and facilitate drug discovery.

A key challenge associated with PDO models is their inability to retain the TME and capture complex interactions among cancer cells and the TME. In our cohort, TME-related genes were downregulated in PDOs compared to tumors, as expected when culturing PDOs independent of their microenvironment^28^. To address this limitation, we are biobanking autologous cancer-associated fibroblasts, tumor-infiltrating immune cells, and peripheral blood mononuclear cells separately, with the aim to develop co-culture systems that integrate distinct components of the TME^28,51^. The ECM also influences gene expression and drug sensitivity, yet current culture matrices like Matrigel are undefined and lack physiological relevance. Using synthetic hydrogel-based matrices could provide a more controlled environment and reveal how matrix composition alters inhibitor responses^52^. We believe these co-cultures and hydrogel-based cultures will enable the study of cancer cell interactions with the TME and provide insights into tumor self-seeding and resistance mechanisms.

Another key challenge for PDOs is the time required to generate stable models. While we prioritized establishing PDOs as long-term models, we acknowledge that tumors evolve over time, and PDOs — especially those requiring longer establishment times — may not fully capture the dynamic landscape of parent tumor. Some platforms address this concern by conducting experiments immediately after dissociating specimens into single cells. This temporarily preserves the diversity of cell types from the parent tissue in short-term cultures^53,54^, but may limit the number of drugs that can be tested.

Despite these challenges, PDOs are valuable models for functional precision oncology, as demonstrated by our assessment of PARPi sensitivity in selected PDOs. Our drug screening confirmed the feasibility of using PDOs as functional precision oncology models for assessing Talazoparib responsiveness. While BRCA2 mutations were linked to sensitivity, many sensitive PDOs lacked known BRCA risk mutations, suggesting that additional mechanism contribute to Talazoparib sensitivity, potentially expanding PARPi indications beyond current biomarkers. We identified alternative molecular signatures, including chromatin remodeling and stress response genes, associated with sensitivity. Notably, distinct BRCA2 variants, such as the pathogenic N3124I and the recurrent S1979R variants (currently classified as VUS), were linked to Talazoparib response. Pathway analysis in sensitive PDOs revealed upregulation of mismatch repair and ECM-related pathways, along with downregulation of antigen-presenting pathways, indicating a complex interplay of DNA repair reliance and immune evasion mechanisms. Additionally, combination drug screening identified several inhibitors that enhanced Talazoparib sensitivity, with the strongest synergy observed with Temozolomide^33^. Synergy often co-occurred with TP53 mutations and an alteration in DNA damage repair pathways. These results highlight the importance for comprehensive molecular profiling and direct drug sensitivity testing with PDO models to enhance precision oncology strategies and expand therapeutic applications.

In summary, we have developed a well-characterized pan-cancer PDO platform that includes histopathology, genomic and transcriptomic comparison with their parent tumors, This resource serves as a valuable tool to advance cancer modeling and support research in functional precision oncology, illustrated by the inclusion of some of our models in the global Human Cancer Model Initiative (HCMI)^55^.

## ACKNOWLEDGMENTS

This work was supported by the Englander Institute for Precision Medicine. Additional project support was provided in part by the Center for Translational Pathology within the Department of Pathology and Laboratory Medicine at Weill Cornell Medicine. This work was also supported by the Department of Defense Prostate Cancer Research Program Health Disparity Research Award PC200267. M.D.G. is supported by 2020 AACR-The Mark Foundation for Cancer Research “Science of the Patient” Grants 20-60-51-GONC, the National Cancer Institute CA279561, and the Manoogian Simone Foundation. J.H.Z, was supported by National Institute of Arthritis and Musculoskeletal and Skin Diseases R01AR077664.

---

**Supplementary Figure 1.** Overview of the PDOs Cohort. Breakdown of the current PDO status **A)** collected over years, **B)** primary tumor location, and **C)** specimen type. **D)** H&E-stained slides from three representative PDO and parent tumor pairs highlight notable morphological differences between the PDOs and their corresponding tumor biopsy sections, underscoring the importance of histopathological review and potential sampling limitations during PDO generation Left: Biopsy from a patient with a history of high-grade urothelial carcinoma showed primarily benign urothelial cells with some reactive changes; however, the corresponding PDO exhibited features of urothelial carcinoma, including marked nuclear pleomorphism and increased proliferation. Middle: Cancerous PDOs were derived from a biopsy that appeared histologically benign endometrial tissue. Right: In contrast, PDOs derived from a biopsy diagnosed as metastatic bladder cancer displayed benign endometrial morphology. Scale bar: 150 μm.

**Supplementary Figure 2.** Correlations and concordance for mutation profiles between PDOs and tumors in the cohort. **A)** Scatter plot with a linear regression line (and gray 95% confidence intervals) between the paired tumor and the PDO ploidies and 95% confidence intervals. X-axis show the PDO ploidy and y-axis show the tumor ploidy. **B)** A bagplot to show the spread, center, and the 21 outliers (marked as red) for the bivariate ploidy dataset which includes the PDO ploidy (x-axis) and tumor ploidy (y-axis). **C)** Scatter plot with a linear regression line (and gray region for 95% confidence intervals) to show no correlation between genomic concordance (x-axis) and tumor purity (y-axis). **D)** Boxplots showing the distribution of genomic concordance for 200 cancer driver genes^18^ between PDOs and parent tumors across 12 cancer types, calculated based on the proportion of shared non-silent mutations from WES. **E)** Boxplots showing the distribution of genomic concordance for genes from targeted panels between PDOs and parent tumors across 11 cancer types. **F)** Oncoprint for mutations in targeted panels grouped by PDO-tumor pairs. The plot shows mutations present at ≥ 5% in both tumor and PDO groups.

**Supplementary Figure 3.** Oncoprints for cancer specific driver gene mutations^18^ in a paired analysis of PDO and parent tumors for **A)** Colorectal, **B)** Lung, **C)** Bladder, **D)** Endometrium, **E)** Prostate, **F)** Pancreatic, **G)** GI, Stomach and Esophageal, **H)** Ovarian cancers.

**Supplementary Figure 4.** Clonal dynamics between paired tumor and PDOs. **A)** Graphical representation of shared clonal architecture between pairs of PDOs and parent tumors from the breast cancer sub-cohort. Each clonal tree is associated to a WCM sample if and genomic concordance shown in brackets. Text color yellow indicates high concordance of ≥ 80% and text in color blue indicates low concordance group of < 40%. Clonal trees are also annotated with known driver mutations in the corresponding samples. Sphere of cells to the left of each tree represent proportion of clonal subpopulations in a sample out of a 100%. **B)** Barplots showing the median percentage (± Inter Quartile Range [IQR]) of dominant tumor clones across different primary tumor sites. Clone status was defined based on changes in cellular prevalence between tumor and PDO: Expanded (> 1.25X increase), Contracted (< 0.75X decrease), Unchanged (within ±25%), or Lost (detected in tumor, absent in PDO). Only clones with > 10% cellular prevalence in the tumor were considered dominant and used for this analysis. Tumor types with fewer than two matched pairs were excluded.

**Supplementary Figure 5.** Comparative analysis of CNA and mutation profiles for PDOs and tumors in the cohort. **A)** Scatter plot to show no correlation between CNA concordance (x-axis) and tumor purity (y-axis). **B)** Scatter plot to show correlation between CNA concordance (x-axis) and difference in fraction genome altered (FGA) values for PDO minus FGAs for parent tumors (y-axis) CNA concordance (x-axis) and tumor purity (y-axis). Both scatter plots have a linear regression line with a gray shaded region for 95% confidence interval. **C)** Oncoprint displaying known driver CNAs in the cohort, split by tumors versus PDOs and ordered by concordance groups: low (< 40%), mid (40-80%), and high (≥ 80%). Barplots on the left show the proportion of matched PDO-tumor pairs that shared the corresponding alteration compared to all pairs within each concordance group. **D)** Oncoprint showing known cancer driver genes and driver CNAs in the PDOs belonging to the low concordance group (< 40%), highlighting the utility of PDO models in cancer research despite low concordance with parent tumors.

**Supplementary Figure 6.** GISTIC analysis of the tumor and PDO samples analyzed as two separate groups. Plots showing significantly **A)** deleted and **B)** amplified CNA peaks identified in the groups of either tumors or PDOs. Left panel shows GISTIC results for PDOs and right panel shows it for tumors. Orange dashed line denotes a GISTIC p-value of < 0.01. **C)** Plots showing a subset of deleted peaks restricted to those with GISTIC p-values of < 0.01 in either PDOs or tumors (but not present in both). **D)** Plots showing a subset of amplified peaks restricted to those with GISTIC p-values of < 0.01 in either PDOs or tumors (but not present in both).

**Supplementary Figure 7.** Cancer-specific DGE analysis. **A)** Volcano plots showing upregulated and downregulated genes identified in each DGE analysis between PDOs from early passages and tumors. DGE was performed for cancer types with at least 3 PDO-tumor matched pairs. Tumors were used as the reference groups in all DGEs. **B)** Bar plots showing the number of DGE genes identified either uniquely in a cancer type or shared among multiple cancer types, as marked by black dots in the upset plot at the bottom.

**Supplementary Figure 8.** Overview of analyses comparing Talazoparib sensitivity in PDO samples. **A)**. Representative dot plot showing changes in PDO area change every 24 hours relative to the day of drug administration, following treatment with 50 µM Olaparib. Each line represents a distinct PDO line. **B)** Representative images of PDO staining with RAD51, pγH2AX, and geminin on functional homologous recombination proficient (HRP) and HRD lines at different timepoints post 10 Gy irradiation. **C-D).** Scatter plots illustrating the relationship between Talazoparib IC50 and the percentage of RAD51-positive cells based on primary tumor site **C)** and prior response to platinum treatment and number of treatments **D).** The dash line indicates the cut off for Talazoparib sensitivity, representing 75% cell death at a concentration of 10 μM. **E)** Oncoprint of select genes in PDOs, grouped by their sensitivity (s) or resistance (r) to Talazoparib. Genes included in the oncoprint either have a Fisher’s Exact Test p-value < 0.05 or a frequency difference greater than 30% between groups, with one group showing zero frequency. **F)** Barplots showing pathways upregulate in sensitive versus resistant groups in a GSEA analysis. **G)** Barplots showing pathways downregulate in sensitive versus resistant groups in a GSEA analysis. Pathways are grouped by their broader biological function.

**Supplementary Figure 9.** Heatmaps for Talazoparib and selected inhibitors generated using the ZIP model in SynergyFinder+. Percent viabilities at 10 μM Talazoparib of each line are included. Red circled box indicates synergistic effects when mean score is above 10.

**Supplementary Table 1.** Summary of the number of PDOs established and characterized by histopathology and genomics.

**Supplementary Table 2.** Summary of minor and major morphological discrepancies between PDO and parent tumors identified in the histopathology review.

**Supplementary Table 3.** Detailed Metadata, Histopathology, Genomics, and Transcriptomics for Established PDO Cohorts

**Supplementary Table 4.** Frequency of 200 pan-cancer driver genes in the cohort^18^.

**Supplementary Table 5.** Genes with non-silent mutations showing differences between PDO and parent tumor Groups (p < 0.05). None of the genes remained significant after false discovery rate (FDR) correction.

**Supplementary Table 6.** Frequency of cancer-specific driver genes in the cohort.

**Supplementary Table 7.** Frequency of cancer specific driver genes by concordance group

**Supplementary Table 8.** Significant CNA regions identified in PDOs or tumors from GISTIC analysis (q value < 0.01)

**Supplementary Table 9.** Number of differentially expressed genes uniquely identified in the given cancer type.

**Supplementary Table 10.** Summary table showing the genomic profiles and drug sensitivity of selected PDOs tested with PARPi.

**Supplementary Table 11.** Media Formula for Each Tumor Type

**Supplementary Table 12.** Catalog number of all materials used in the paper.

**Supplementary Table 13.** Cancer specific antibody used in IHC for pathology review.

## METHODS

### Sample Collection

The tumor organoid platform at the Englander Institute for Precision Medicine develops PDOs through biopsies, surgical resections, or rapid autopsy procedures. Fresh tissue samples were collected with written informed patient consent with the approval from Weill Cornell Medicine (WCM) IRBs: #0908010582, #1807019405, #1008011221, #1011011386, #19-05020075, #1305013903, and Memorial Sloan Kettering Cancer Center IRBs: #15-318, #06-107.

### PDO Establishment and Culture

PDOs were established using modified standard protocols^1^. Fresh tissue specimens were collected with portions frozen in Tissue-Tek optimal cutting temperature (OCT) compound for genomic or histopathological analysis. Media formulations are detailed in Supplementary Table 11 and material catalog number are in Supplementary Table 12. Remaining tissue was transported to the laboratory, washed 3 times in Transport Media, mechanically dissected to approximately 2 mm pieces, and enzymatically digested in a 20 times volume of Collagenase Media on a shaker at 200 rpm at 37°C until the solution turned cloudy (30-45 minutes). Resulting cell pellet was centrifuged at 1300 rpm for 3 min, washed with Washing Media, and resuspended in a cell media/Matrigel mixture (v/v 1:2) with tissue-specific culture media. Up to five 100 μL drops of drops were plated into a 6-well cell suspension culture plate, polymerized at 37°C for 30 min, and overlaid with 3 mL tissue-specific culture media. Media was refreshed every 3-4 days. PDO formation is characterized by the observation of a 3D structure, with evidence of cell proliferation and cluster formation. PDOs were passaged in Matrigel using TrypLE Express for 7-10 minutes or Accutase for 5-7 minutes, depending on tumor type, and replated as described above. Monthly mycoplasma testing was performed using the PCR Mycoplasma Detection Kit on cell lysates instead of media. Biobanking was initiated when cell pellets exceeded 1 million cells, using Recovery Cell Culture Freezing Medium for overnight storage in −80°C, followed by transfer to liquid nitrogen.

### PDO Media

Media formulations are detailed in Supplementary Table 11. For homemade Noggin and R-spondin media, HEK-T293 cells with respective cassettes were thawed into a T175 flask with DMEM media, 10% FBS, and P/S. Once cells reach 90-100% confluence, they were split with TrypLE Express and replate into twenty 150mm plates with 25 mL of DMEM media. After 3 days, when cells should have reached at least 70% confluence, cells were starved by aspirating the media, washing with 5 mL PBS, and adding 25 mL of Washing Media. After 24 hours, the conditioned media was collected, filtered through 0.22 uM filters, tested for mycoplasma, and stored at −20°C until use.

### PDO Pathological Characterization

PDO histopathology was verified by comparing fixed passage 5 organoid sections to parent tumor sections using our cytology and histology platforms^2,3^. PDOs were released from Matrigel droplets using Cell Recovery Solution, suspended in a Fibrinogen/Thrombin (v/v 10:1) pellet, and fixed with 4% paraformaldehyde in PBS. Samples were then sent to WCM Center for Translational Pathology for formalin fixation, paraffin embedding, and staining. H&E, Ki67, and additional cancer-specific staining are listed in Supplemental Table 14. A comprehensive cytomorphologic comparison of PDTO and primary lung tumor (Histopathology H&E section) was performed by evaluating cytological features, including nuclear and cytoplasmic characteristics, as well as morphological features. Additionally, IHC was conducted and compared between the tumor and the organoid where necessary.

### DNA and RNA Extraction

Genomic DNA and RNA were extracted from various samples, including formalin-fixed paraffin-embedded (FFPE) tissues, OCT-embedded frozen tissue, PBMCs, saliva, and organoid cell pellets for profiling. Material catalog number are in Supplementary Table 12. DNA from FFPE tissue was extracted using the Maxwell 16 FFPE Plus DNA kit on the Maxwell 16 instrument. Tissue sections (10 µm) were macro-dissected based on the annotated H&E slides. DNA from OCT-embedded tissue was extracted using Maxwell 16 Tissue DNA Purification Kit or Omega Bio-Tek Mag-Bind Blood & Tissue DNA HDQ kit. For saliva, DNA was extracted using Oragene collection kit. DNA from buffy coat fractions was extracted using Maxwell® 16 LEV Blood DNA Kit or Omega Bio-Tek Mag-Bind® Blood & Tissue DNA HDQ kit. Organoid cell DNA was extracted using Qiagen DNeasy Blood & Tissue Kit or Omega-Bio-Tek Mag-Bind Blood & Tissue DNA HDQ kit. RNA from organoid cells was extracted with Maxwell 16 LEV simplyRNA Cells kit. DNA quantity and quality were assessed using a Qubit 4 Fluorometer and an Agilent Tapestation 4200 machine. RNA was similarly quantified using a Qubit 4 Fluorometer, and quality evaluated using an Agilent 2100 Bioanalyzer.

### Whole Exome Sequencing (WES)

#### WES Library Preparation and Sequencing

The Exome Cancer Test v1.0 (EXaCT-1) library preparation was done using FFPE DNA as templates for PCR amplification of two independent GAPDH amplicons^4^. Amplicon yields from FFPE DNA were measured with Agilent Bioanalyzer and compared to those from intact reference DNA template. The sample-to-reference yield ratio served as a quantitative indicator of FFPE DNA integrity, predicting sample performance in HaloPlex target enrichment. FFPE DNA samples were categorized based on yield ratio: A (>0.2), B (0.05∼0.2) and C (<0.05), with input DNA amounts adjusted accordingly. Acceptable DNA requirements included >225 ng for fresh frozen high molecular weight DNA, >500ng for high-quality FFPE DNA, and >1000ng for low-quality FFPE DNA, with Qubit 4 Fluorometer absorbance ratios of A260/A280 (1.8∼2.0), A260/A230 (>2.0).

For the samples processed after the end of the year 2020, a newer version of the Exome Cancer Test, called EXaCT-2, was used. EXaCT-2 is based on SureSelect Human All Exon V6^5^, with augmented coverage of approximately 1400 cancer-related genes, enabling the detection of mutations at very low allelic frequencies. The capture kit also included chromosome tiling probes in intergenic regions to provide better analysis of copy number alterations^6^. Less than 200 ng of FFPE DNA sample was enzymatically fragmented using SureSelectXT HS and XT Low Input enzymatic fragmentation kit. Material catalog number are in Supplementary Table 12. The fragmented DNA underwent end repair, A-tailing, and adapter ligation with SureSelect XT Low Input Reagent Kit. Each library was made with a unique dual index with SureSelect XT Low Input Dual Index P5 Indexed Adaptors. Then, the indexed libraries were captured using the custom-designed EXaCT-2 Target Enrichment Human All Exon probes, amplified, and their quality and quantities were assessed by Agilent 4200 TapeStation System and Invitrogen Qubit Fluorometer. Final libraries were pooled and sequenced on an Illumina NovaSeq6000 sequencer at PE 2×150 cycles.

#### WES Data Alignment and Processing

Raw sequencing reads in BCL format were processed through bcl2fastq 2.19 (Illumina) for FASTQ conversion and demultiplexing. Quality control was performed with fastp^7^. The sequencing reads were aligned to the human reference genome *hs37d5* using the Burrows–Wheeler Alignment tool (bwa v0.7.17, using ‘mem’ option). After alignment, PCR duplicates were marked using Sambamba (v0.6.7) for samples sequenced with SureSelect. Samples sequenced with HaloPlex did not require this step since HaloPlex is amplicon-based. This was followed by realignment around the indels, fixing mates and base quality score recalibration using Genome Analysis Toolkit (GATK, v4.2.0.0), and all unmapped reads (MAPQ = 0) being removed. Accurate pairing between each tumor (or PDO) and its corresponding germline was determined using SPIA (Single Nucleotide Polymorphism [SNP] Panel Identification Assay)^8^, which calculates genetic distance between samples based on a selected set of SNPs. Sample pairs with a SPIA score of > 0.4 were deemed mismatched and excluded from further genomic analysis. CLONET^9^ was applied to estimate tumor purity and ploidy. MSI for each sample was calculated using MSIsensor-pro^10^.

#### Somatic Mutation Detection

For EXaCT-1, somatic mutations were identified in matched tumor (or PDO) and normal pairs using SNVseeqer^11^. Mutations were filtered based on a minimum read depth of 10 in the tumor or PDO with at least 2 reads supporting the alternate allele. Variant allele frequency (VAF) filters were applied as follows: ≥ 25% for variants of unknown significance and ≥ 10% for known cancer genes based on COSMIC Cancer Census^12^. A list of clinically relevant mutations based on the EIPM Knowledge Base was always reported. The final set of mutations were annotated using Oncotator^13^.

For EXaCT-2, somatic mutations were identified using a method adapted from the Multi-Center Mutation Calling in Multiple Cancers (MC3) pipeline^14^. The adapted pipeline included six SNV callers (VarScan^15^, SomaticSniper^16^, MuTect2^17^, Strelka2^18^, RADIA^19^, and MuSE^20^) and four indel callers (VarScan^15^, MuTect2^17^, Strelka2^18^, Pindel^21^). Each tool was run independently, and all mutations identified were merged. When combining overlapping mutations, the read depth and VAFs from different tools were averaged. This combined set of variants was then filtered based on the following criteria: within-sample allele strand bias frequency (0.1), a total read depth of 30X in tumor or PDO, and a population frequency greater than 0.01 in either TOPmed (Trans-Omics for Precision Medicine^22^) or gnomAD (Genome Aggregation Database^23^). Frequently mutated, but biologically unrelated genes such as *TTN* and the *MUC* were also excluded^6^. The final set of mutations from EXaCT-2 were annotated using Nirvana (v3.3.103.11)^24^. Tumor Mutation Burden (TMB) was calculated for each sample by counting the total number of non-synonymous mutations (including truncating mutations), normalized to the number of bases in the targeted exonic regions per million. Mutations that correlated differently with one group versus the other were identified using Fisher’s Exact Test and a p-value threshold of < 0.05.

#### Copy-number Analysis

Copy-number alterations (CNV) were identified from aligned reads using CNVkit v0.9.10^25^. Segments with a log2 ratio of ≥ 0.2 were classified as copy number gains, while those with a log2 ratio of ≤ −0.2 were classified as copy number losses. These copy number calls were then adjusted for tumor purity and ploidy using CLONET v2^9^. For gene-level analysis, copy number segments were annotated for genes based on GENCODE v19^26^. Log2 ratios for multiple segments spanning a gene were combined by calculating a weighted average based on the overlap of segment coordinates, resulting in a gene-specific log2 ratio. The Fraction of Genome Altered (FGA) was calculated by dividing the total number of bases with copy number gains or losses by the size of the human genome (3,054,815,472 bases).

Driver CNVs were identified using GISTIC2 (v2.0.23)^27^ applied to the segmented log2 ratios. GISTIC was run with the reference file *hg19.UCSC.add_miR.140312.refgene.mat* and a confidence threshold of 0.99. This analysis was conducted separately for tumors and PDOs in the cohort. CNV peaks that were differentially deleted or amplified in either tumor or PDO, but not in both, were identified based on a p-value of < 0.01. Network analysis of genes within these significant CNV peaks was performed using the *enrichGO* function from the *clusterProfiler* package *(v4.6.2)* visualized with the *cnetplot* function from the *enrichplot* package (v1.22.0) in R.

#### Identification of Mutational Signatures

Mutational Signature analysis was performed using deconstructSigs (v 1.9.0)^28^. First, the *mut.to.sigs.input* function was used to transform the mutation data into a matrix of 96 possible trinucleotide contexts. Then, the *which Signatures* function was applied to infer the contributions of each sample to the 60 COSMIC signatures from the reference *signatures.exome.cosmic.v3.may2019*, with the *tri.counts.method* parameter set to “exome”. To evaluate the similarity between mutational signature profiles of tumor and PDO pairs, cosine similarity values were computed using the *lsa* package (v0.73.3) in R. The contributions of the most frequent signatures in the cohort were visualized using the *ComplexHeatmap* package (v2.13.0) in R.

### Targeted Gene Panels

#### Library Preparation and Sequencing for Targeted Panels

Our targeted gene panels included the TruSight Oncology 500 (TSO500; Illumina) and the Oncomine Focus Assay (ThermoFisher Scientific). Library preparation and sequencing were performed by WCM Clinical Genomics Laboratory. DNA was extracted from FFPE or cell pellets with a minimal concentration of 3.33 ng/uL using Maxwell® 16 FFPE Plus DNA kit. Material catalog number are in Supplementary Table 12. Genomic DNA was fragmented into 90-250 bp using Covaris E220 evolution Focused-ultrasonicator. For TSO500 sequencing, TSO500 DNA/RNA High-Throughput Kit was used for end repair, A-tailing, and adapter ligation. Libraries were amplified, indexed with Illumina DNA/RNA UD indexes, and captured through dual hybridization, then enriched and cleaned up. Libraries were normalized using the TSO500 bead-based normalization method, and quantity was checked using Qubit dsDNA HS Assay Kit. Sequencing was performed on NovaSeq 6000 SP Reagent Kit and NovaSeq6000 system (Illumina) with PhiX Control v3 as control. Analysis was performed using Illumina TruSight 500 Local App v2.2. For Oncomine, DNA targets were amplified with Ion AmpliSeq Library Kit with DNA primer pools. Primer sequences were partially digested, ligated to the Ion Xpress Barcode Adaptors, and purified with Agencourt AMPure XP Kit. The library was quantified by comparing it to *Escherichia coli* DH10B control library with duplicates showing SD values higher than 5.83 repeated. Sample libraries were diluted to 50 pM for sequencing. Samples were loaded on Ion 540 Chip Kitand Ion 540™ Kit-Chef.

#### Variant Detection in Targeted Panels

For TSO500, the sequenced reads were analyzed using the TruSight Oncology 500 LocalApp 2.2 pipeline and Pindel (v0.2.5b9, 20160729)^21^. Indels were filtered to include variants larger than 20bps with at least 10X depth that did not appear in population databases at > 1%. Pathologists manually reviewed clinically relevant hotspot variants that might be missed by the pipeline. The filtered variants were reviewed using Velsera’s Clinical Genomics Workspace for clinical annotation. For Oncomine, the sequenced reads were analyzed using the standard Oncomine Comprehensive Assay v2 (OCAv2, ThermoFisher Scientific) Ion Reporter software (v5.0) with “OnCORseq Comprehensive v2.0 - DNA - Single Sample” workflow (v5.6), previously validated at WCM^29^. To better capture complex indels, an additional workflow was run with SNP realignment disabled and the “*Allow_Complex*” setting enabled. Identified variants were annotated using the Torrent Variant Annotator v2.0 plug-in. Synonymous variants occurring at a frequency >1% in the 5000 Exomes or ExAC databases (within Ion Reporter) were excluded. The identified variants were manually reviewed by pathologists and summarized in clinical reports.

### RNA Sequencing (RNA-Seq)

#### RNA-Seq Library Preparation and Sequencing

RNA libraries preparation and sequencing was performed by WCM Genomics Core Laboratory. For polyA selection, samples were processed using Illumina TruSeq Stranded mRNA Sample Library Preparation kit and Agilent high throughput sample preparation Bravo B system according to the manufacturer’s instructions. Material catalog number are in Supplementary Table 12. Normalized cDNA libraries were pooled and sequenced on Illumina NovaSeq 6000 sequencer with pair-end 100 cycles. Raw sequencing reads were converted from BCL to FASTQ format using bcl2fastq 2.19 (Illumina). For rRNA depletion method, rRNA was removed from Total RNA using Illumina Ribo Zero Gold for human/mouse/rat kit. mRNA was fragmented, reverse-transcribed into cDNA using Illumina TruSeq Stranded Total RNA Sample Library Preparation kit. The cDNA fragments underwent end repair, adapter ligation, and PCR enrichment to create the final cDNA library. These cDNA libraries were pooled and sequenced on Illumina NovaSeq 6000 sequencer with pair-end 100 cycles.

#### RNA-Seq Alignment and Quantification

RNA sequencing analysis was conducted as previously described^30^. Sequenced reads were mapped to the human reference genome hg19/GRCh37 using the STAR aligner (RRID:SCR_004463) v2.7.2b and STAR-Fusion version 1.9.1^31^. HTSeq v0.11.3^32^ (RRID:SCR_005514) was used to quantify gene counts and cufflinks v2.2.1^33^ (RRID:SCR_014597) was used to quantify Fragments Per Kilobase of transcript per Million mapped reads (FPKMs) with GENCODE v19 (RRID:SCR_014966)^34^. Tumor purity was inferred from FPKMs using “Estimation of STromal and Immune cells in MAlignant Tumours using Expression data” (ESTIMATE)^35^.

#### Non-supervised Clustering of Expression Profiles

Non-supervised clustering was performed using t-distributed Stochastic Neighbor Embedding (tSNE) on normalized counts from all PDOs. The tSNE algorithm was applied to reduce the dimensionality of the data while preserving the local structure of the expression patterns, allowing for the visualization of clusters based on gene expression similarity. The R tsne package (v0.17) in R was used to implement the tSNE algorithm. The resulting tSNE plot was used to assess the clustering of PDOs by their cancer types.

#### Cancer Hallmark Pathways Analysis

Hallmark pathway Enrichment Scores (ES) were calculated for each sample using single-sample Gene Set Enrichment Analysis (ssGSEA)^36^, implemented with GSVA package (version 1.46.0) in R (version 4.2.2). The signature set used for this analysis was obtained from the Human Molecular Signatures Database (MSigDB) collection of 50 hallmark pathways (H)^37^. Linear regression models were applied using the *lm* function in R (version 4.2.2) to identify differentially enriched hallmark pathways between PDOs and tumors within each cancer type. The linear model equation was as follows: ES ∼ TO + Pair + TP, where ES represents enrichment scores for a given hallmark pathway, TO is a binary variable (0 for tumor, 1 for PDO), Pair denotes factor covariates representing PDO-tumor pairs, and TP stands for tumor purity, inferred from ESTIMATE. Beta coefficients (β) from the linear model were converted to odds ratio (OR) using the formula 10^β^. P-values from all models were adjusted using Bonferroni correction. Differentially enriched pathways were identified using cutoffs of an adjusted p-value < 0.1 and an absolute OR of > 1.5. For visualization, the OR for non-significant pathways was set to 0. The ORs for each hallmark pathway across cancer types were clustered and visualized using the *ComplexHeatmap* package (v2.14.0) in R.

#### Differential Gene Expression Analysis

Differential gene expression (DGE) analysis was performed on counts data using DeSeq2 (v1.42.1)^38^, with PDO-parent tumor pairing and tumor purity (inferred from ESTIMATE) included as covariates. DGE analysis between tumor and early stage PDOs was conducted separately for each cancer type with at least three unique tumor-PDO pairs (n = 7 cancer types). Differentially expressed genes (DEGs) were identified based on an absolute log2 fold change (log2FC) > 2, and an adjusted p-value < 0.01. The results were visualized as volcano plots for each cancer type using the EnhancedVolcano package (v1.20.0). DEGs identified as upregulated (log2FC of > 2) or downregulated (log2FC of < −2) in ≥ 5 cancer types were selected for pathway enrichment analysis using EnrichR^39^. Significantly enriched pathways and biological processes were plotted using the ggpubr package (v0.6.0) in R. To confirm that a DEG was truly unique to a specific cancer type, the DEG was cross-checked against the remaining cancer types using a more lenient threshold of absolute log2FC > 1, adjusted p-value < 0.05, and a base mean > 10 to ensure it wasn’t present in other cancer types. For the DGE analysis of early versus late passage PDOs, all cancer types were analyzed together in a single DESeq2 model, adjusted for primary site.

### Genomic and Transcriptomic Concordance Analysis

#### Somatic Mutations Concordance

Concordance for somatic mutations was assessed for non-silent Single Nucleotide Variants (SNVs) and truncating mutations. SNV concordance was calculated by dividing the number of shared mutations between the PDO and the parent tumor by the total number of mutations in the parent tumor. To minimize the possibility of false non-overlapping mutations, we gathered all mutations from each PDO-tumor pair and manually reviewed them for evidence in samples where the mutation was absent. This review was conducted using bcftools (v1.9) and/or Integrative Genomics Viewer (IGV) (2.3.57). For driver gene SNV concordance, the mutation lists were filtered to include 200 pan-cancer driver genes^40^, and concordance was calculated as described above.

#### Copy-number Concordance

To assess copy-number variant (CNV) concordance, the total number of bases covered by segments in the parent tumor was first calculated, denoted as *x*. Bedtools (v2.28.0) was used to identify overlapping CNV segments between the tumor and the matched PDO. For regions with overlap, the number of bases where the PDO’s copy number status (gain, loss, or neutral) matched that of the tumor was counted, denoted as *y*. CNV concordance was then calculated as y divided by x. Gains were defined as log2 ratios > 0.50, losses as log2 ratios < −0.50, and regions with log2 ratios between −0.50 and 0.50 were considered copy number neutral.

#### Expression Profile Concordance

Expression concordance between a parent tumor and its PDO was calculated using Spearman correlation across top 5000 most variable genes^41^. These correlations were compared to a background distribution generated by randomly pairing PDOs and tumors from the full cohort, ensuring that the true pairs were excluded. Welch Two Sample t-test was used to statistically test if the true and randomly paired distributions were significantly different.

### Clonal Evolution Analysis

All non-silent SNV and truncating mutations, copy number values and tumor purities (inferred from DNA) were used for the clonality analysis. Pyclone VI^42^ was applied to these datasets to infer the clonal clustering of mutations in a paired analysis of parent tumors and PDOs. For genomic locations where mutations were not detected in a sample within the pair, read depths were extracted using bcftools (v1.9). Paired clonal clusters and their corresponding cancer cell fractions (CCFs), along with the VAFs of these mutations in tumor and PDO pairs, were analyzed to understand clonal evolution through clonal ordering and tree visualizations using ClonEvol^43^.

For cohort level analysis, PyClone-VI was run cohort-wide (n=135 pairs) using somatic SNVs, with fixed copy number settings (major allele = 2, minor allele = 0) due to incomplete CNA data^42^. Tumor purity estimates were incorporated where available; for a total of 9 tumor-PDO pairs (5 PDOs and 7 tumors) purity data was not available, for these we assumed tumor purity to 100% for analysis. PyClone ran successfully on 132 tumor-PDO pairs. We treated each mutation cluster identified by PyClone-VI as a putative clone, based on the assumption that variants grouped by similar cellular prevalence represent a shared clonal lineage. To evaluate clonal divergence, we calculated the absolute difference in cellular prevalence between tumor and PDO for each cluster and categorized clusters as Highly Concordant (<10% difference), Moderately Concordant (10%-29%), or Divergent (≥30%). For each tumor–PDO pair, we computed the proportion of clusters in each category and stratified results by SNV concordance status. To assess the fate of dominant clones, we identified clusters with >10% cellular prevalence in the tumor and evaluated their representation in the matched PDO. We calculated fold change between PDO and tumor cellular prevalence to quantify expansion, contraction or loss of dominant tumor clones in PDOs. Clones were labeled as Expanded (>1.25X increase in PDO), Contracted (<0.75X), Unchanged (0.75– 1.25X), or Lost (undetected in PDO).

### PDTO Area Tracking with Incucyte

1,000 cells were plated in a 10-μL cell culture media/Matrigel mix (v/v 1:2) in a 96-well plate. Material catalog number are in Supplementary Table 12. After polymerization at 37°C for 30 minutes, 15 μL of tissue-specific media was added per well. Drugs were introduced 72 hours later in 15 μL of fresh media (total 30 μL) using a 9-point dilution series. PDO growth was monitored via daily brightfield imaging with the Sartorius Incucyte S3, and cell area was analyzed using the affiliated software. All wells were normalized to PDO size to assess drug response.

### PDTO Staining and Microscopy

5,000 cells were plated in 1% Collagen I-coated, black 96-well plates and incubated for 48 hours. Material catalog number are in Supplementary Table 12. Cells were then irradiated at 10 Gy using the RS 2000 ray Irradiator (Rad Source Technologies) and fixed with 4% paraformaldehyde in PBS. Cells were permeabilized and blocked in 0.1% Triton X-100, 2% BSA in PBS for 30 minutes, followed by a 2-hour incubation at 1:400 dilution with antibodies against RAD51, phospho-Histone H2A.X, or geminin. After washing, cells were incubated at 1:500 dilution with Alexa Fluor-conjugated secondary antibodies for 1 hour, then mounted with ProLong Diamond Antifade with DAPI. Images were acquired on a Zeiss LSM 880 confocal microscope (63× objective), with exposure settings adjusted to prevent oversaturation. At least 100 nuclei per condition were quantified, and foci were counted using Fiji ImageJ (v2.14.0/1.54f).

### PDTO Drug Screening

1,200 cells were plated in a 8-μL cell culture media/Matrigel mix (v/v 1:2) in a 384-well plate. Material catalog number are in Supplementary Table 12. After a 30-minute incubation at 37°C for polymerization, 15 uL of tissue-specific media was added per well. Drugs were added 72 hours later in 15 μL of fresh media (total volume: 30 ul) and incubated for 168 hours. Drugs from MedChemExpress or SelleckChem and were diluted per manufacturers’ instructions. Luminescent viability was assessed using CellTiter-Glo 3D, and luminescence was read using the Biotek Synergy Neo2 Plate reader. Data analysis was performed running nonlinear regression in Graph Pad Prism 9. As the reported Cmax for Talazoparib was too low to induce sensitivity^44^, viability at the maximum dose (10 µM) was used to classify sensitivity: viability >25% as resistant and ≤25% as sensitive. Synergy scores were calculated using SynergyFinder+ based on the Zero Interaction Potency (ZIP) model, with scores >10 indicating synergy, <-10 indicating antagonism, and between −10 and 10 suggesting an additive effect.

### Germline Mutation Identification for PARP inhibitor-treated PDOs

Germline mutation in the PDOs used in the PARP inhibitor screening were identified through various methods, including clinical sequencing reports, Exact-1 pipeline, and whole genome sequencing (WGS). Clinical sequencing reports were generated by platforms such as Ambry, MSK-IMPACT, GUARDIAN360, 50 Gene, and Foundation One. Germline somatic variants identified through WES were derived from blood or saliva samples, following the pipeline as previously described^45^. Germline data from selected WGS analysis were obtained from samples included in a separate study^46^.

