## Supplementary Figures for "Pan-Cancer PDOs Preserve Tumor Heterogeneity and Uncover Therapeutic Vulnerabilities"

#### Suppl Figure S1

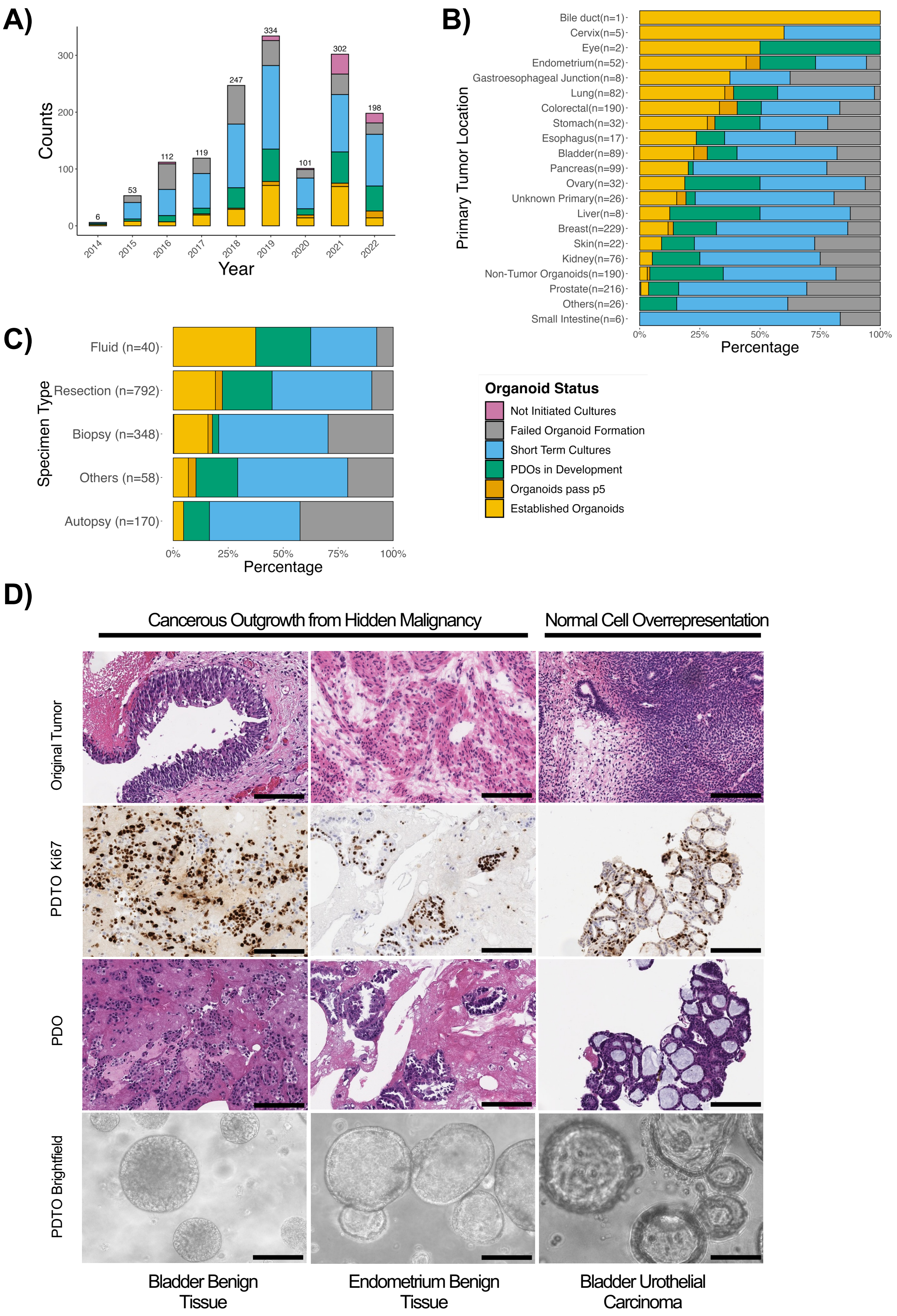

#### Suppl Figure S2

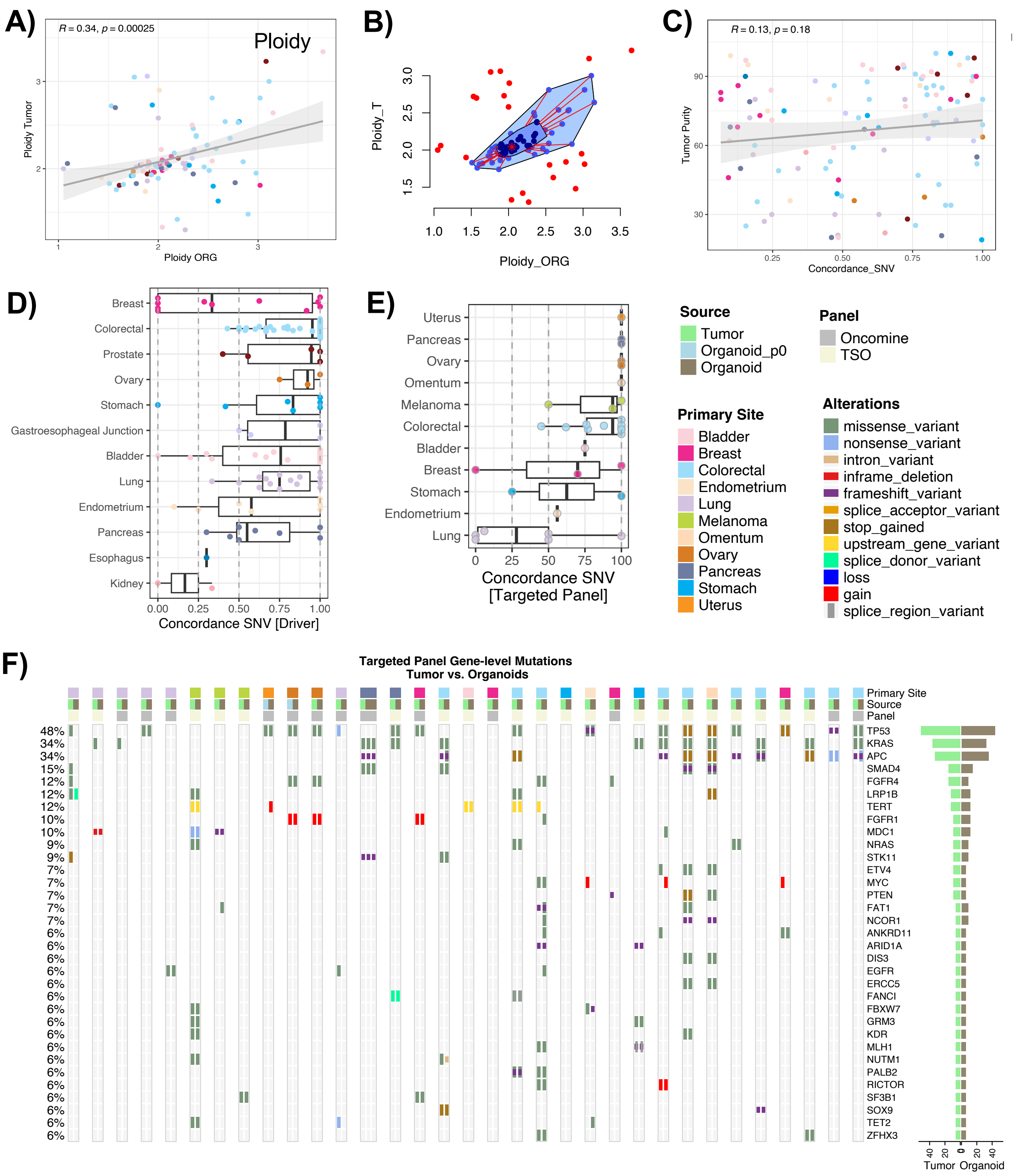

### Suppl Figure S3

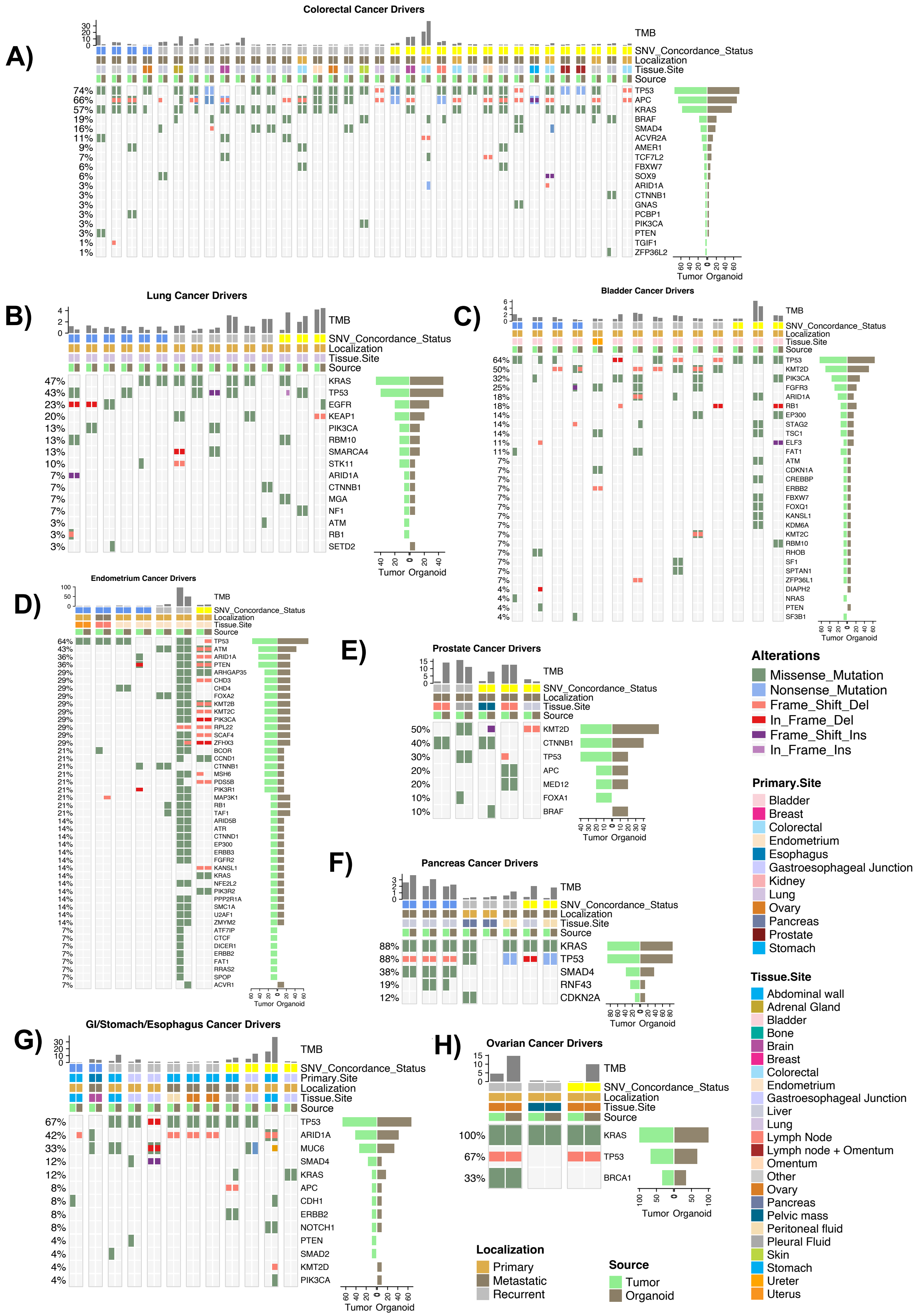

#### Suppl Figure S4

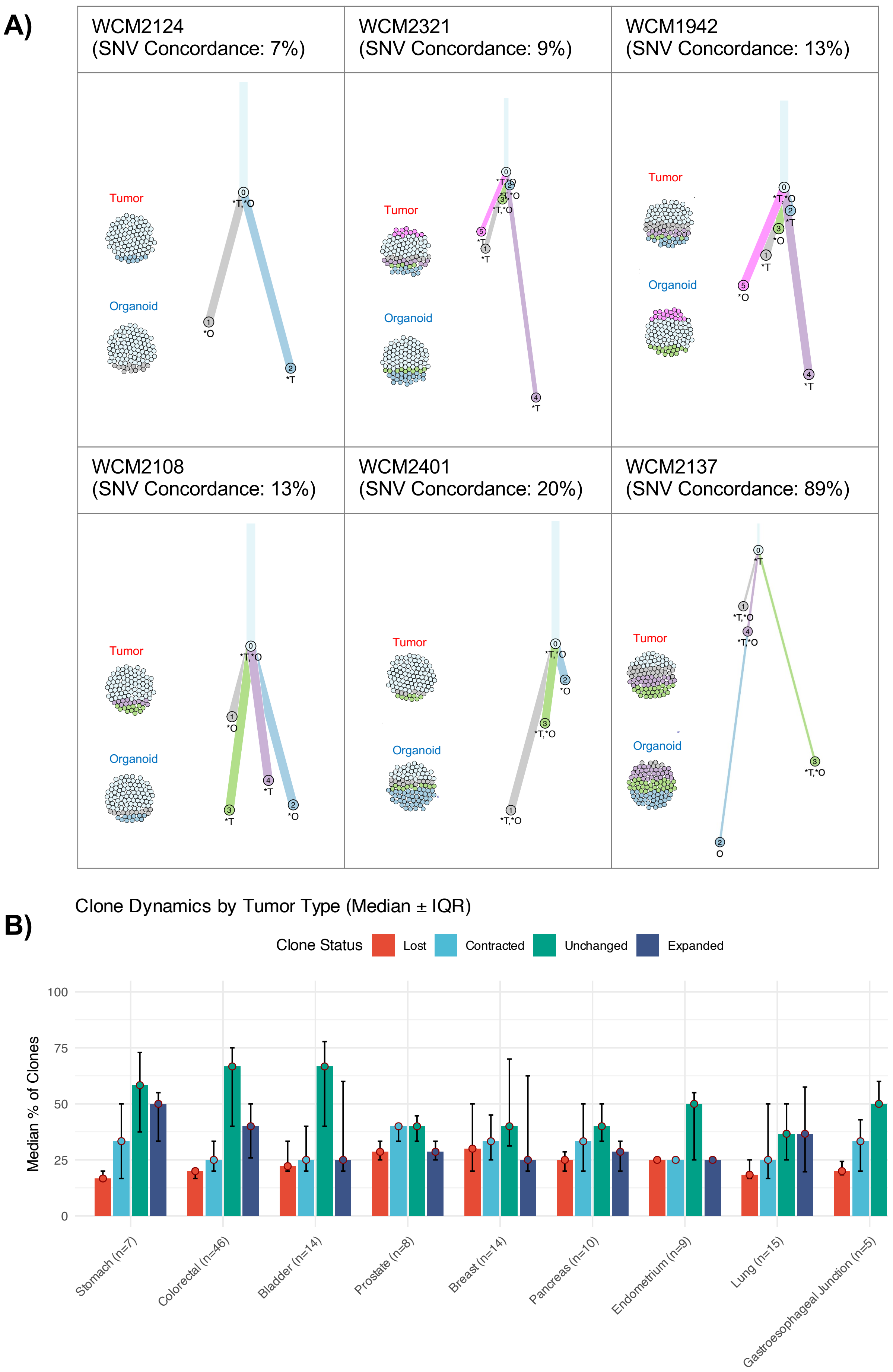

#### Suppl Figure S5

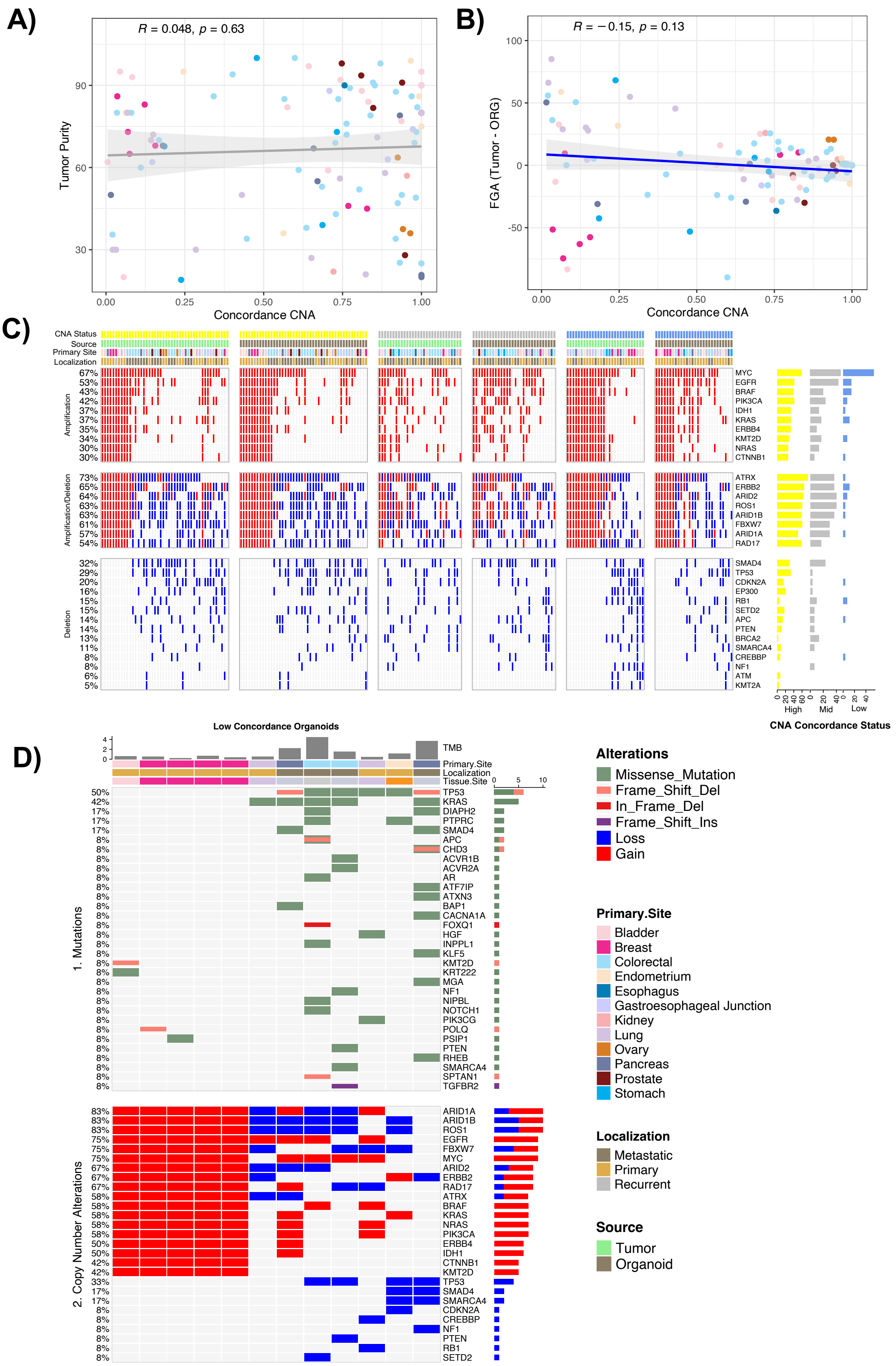

#### Suppl Figure S6

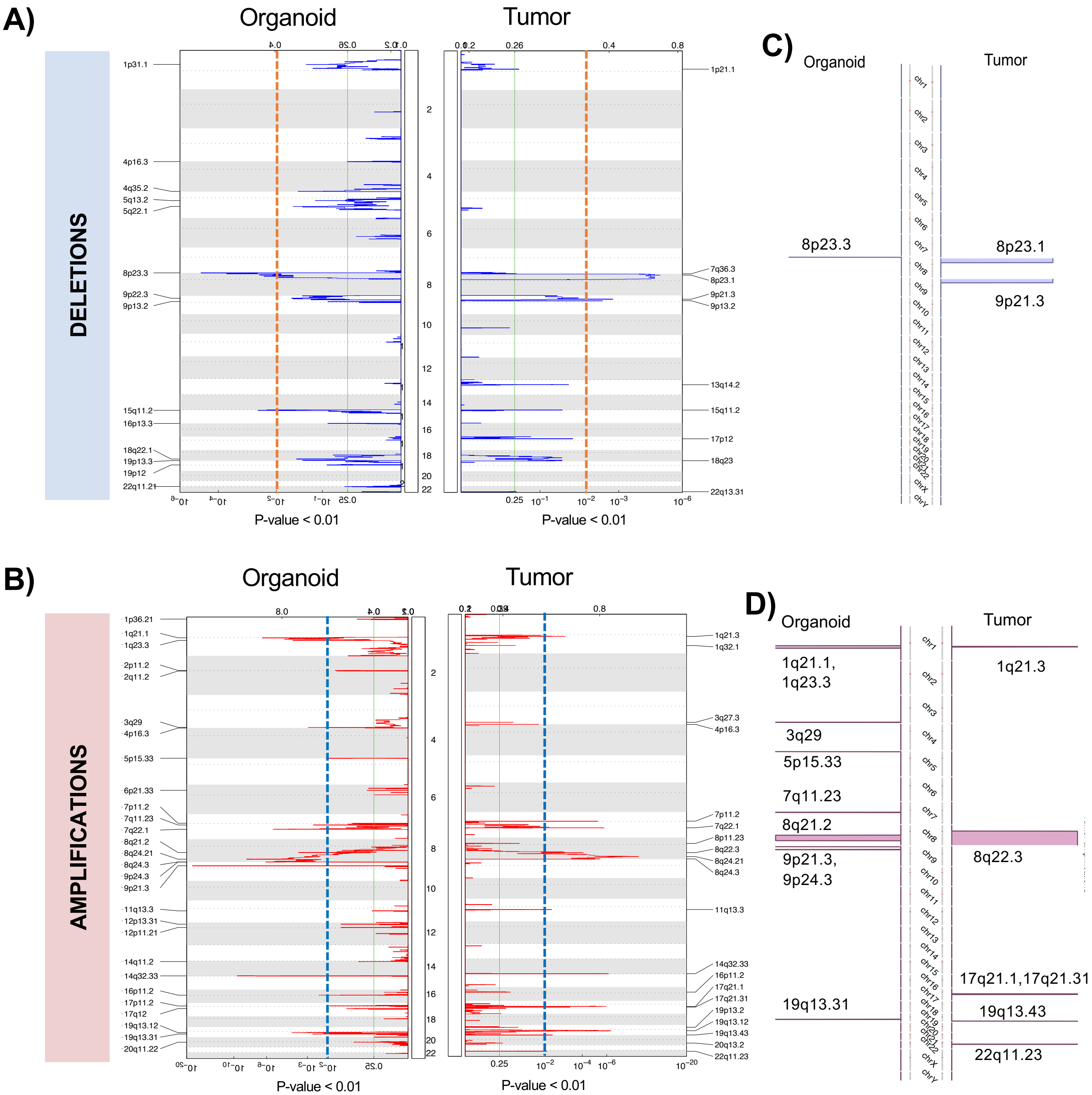

### Suppl Figure S7

A)

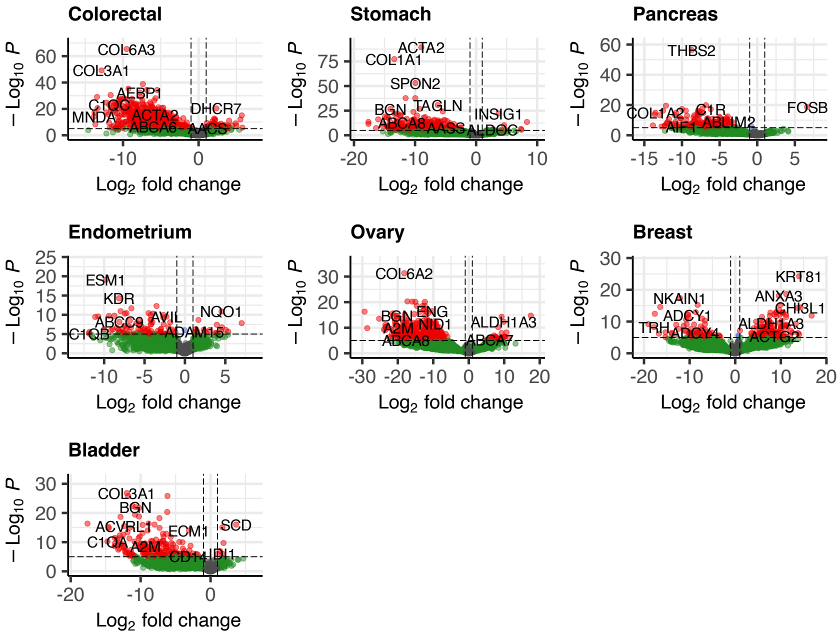


B)

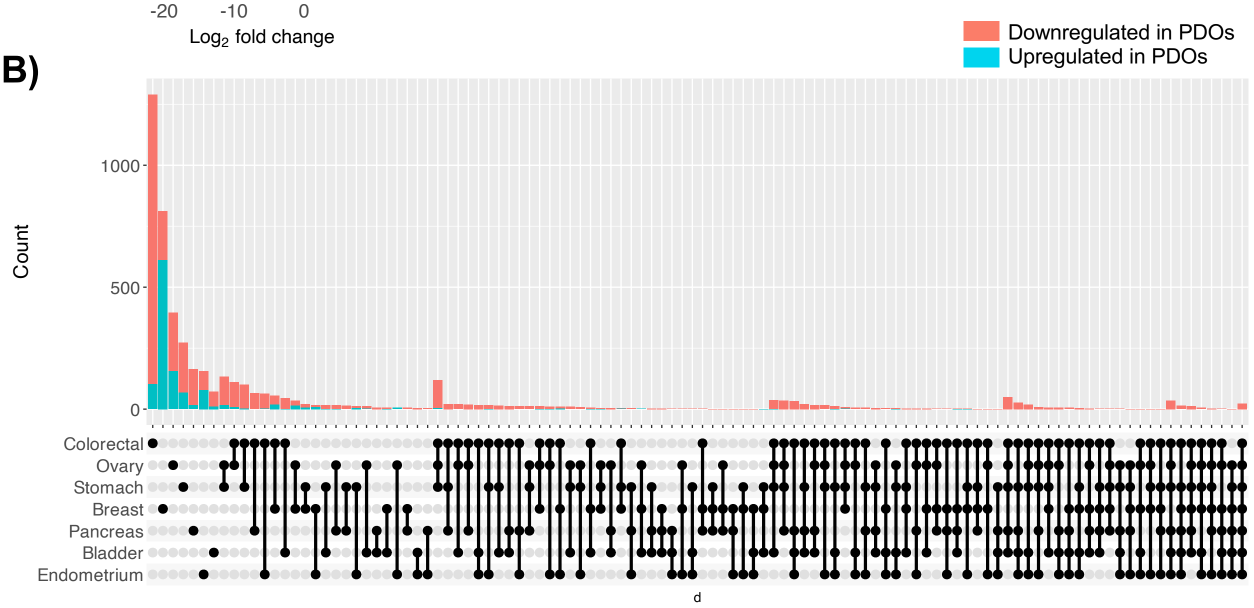

#### Suppl Figure S8

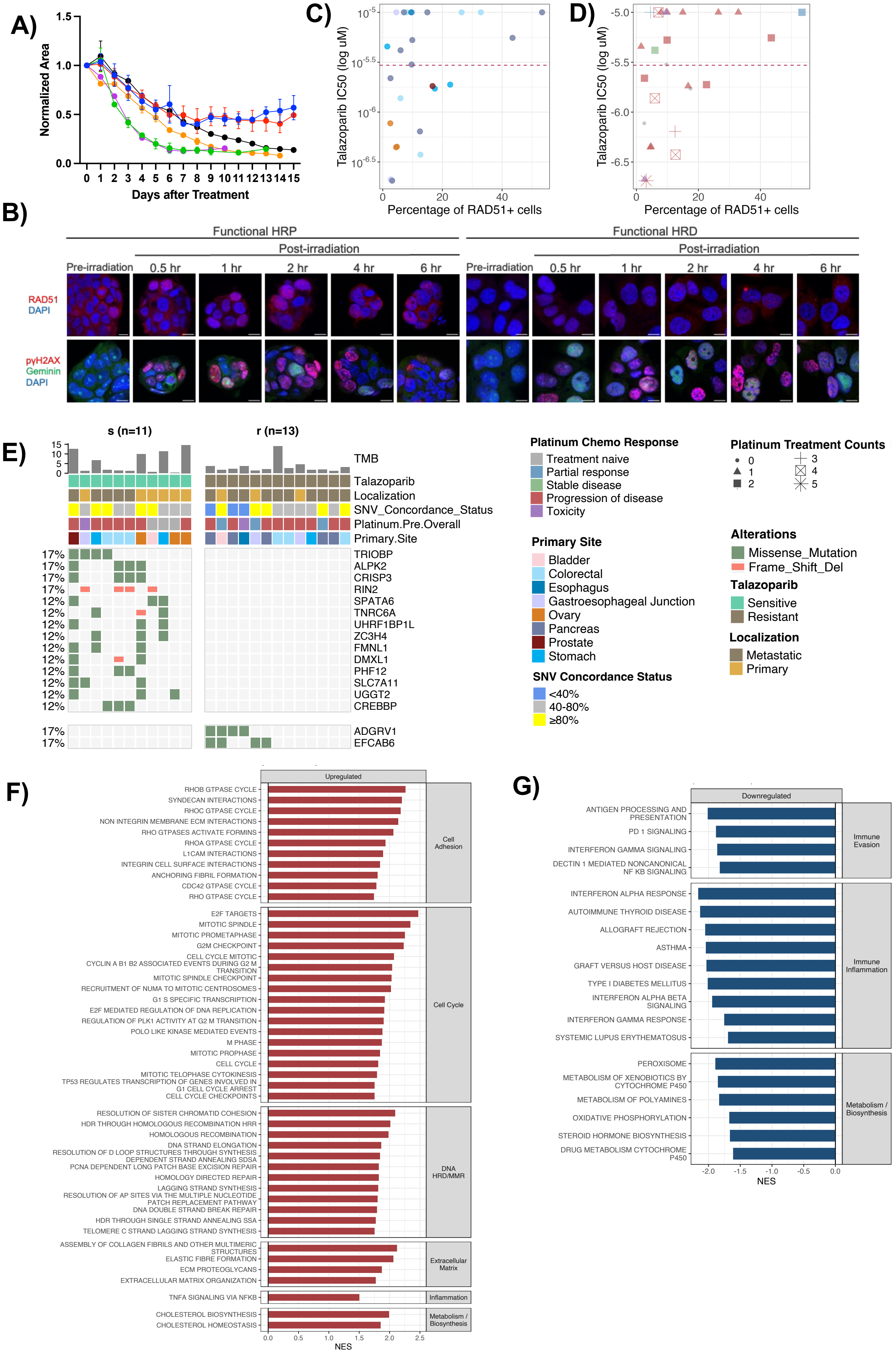

### Suppl Figure S9

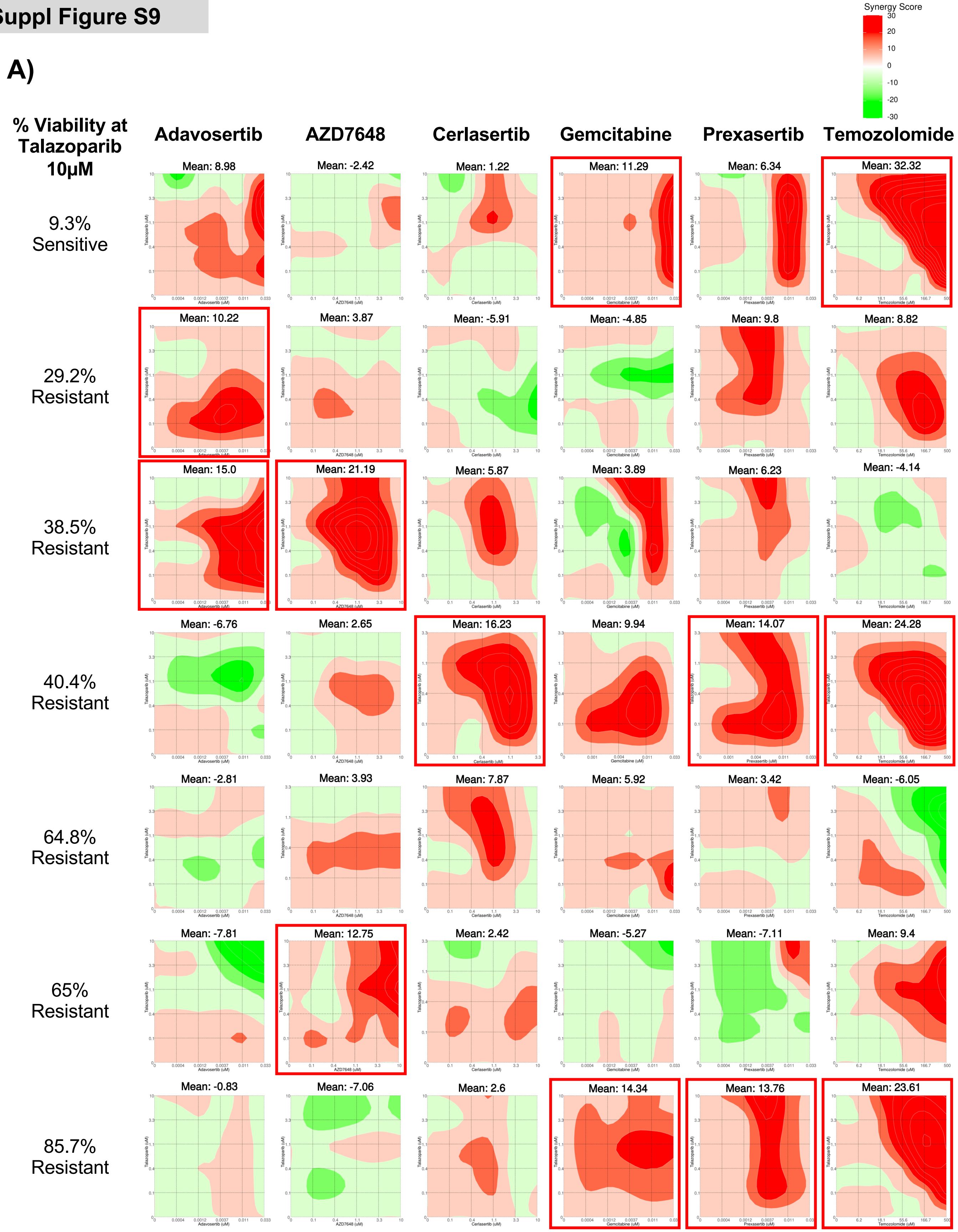
